# Diptera flight diversity is shaped by aerodynamic constraints, scaling, and evolutionary trade-offs

**DOI:** 10.1101/2025.10.20.683381

**Authors:** Camille Le Roy, Ilam Bharathi, Thomas Engels, Florian T. Muijres

## Abstract

Flight has been a key innovation in insect evolution, yet the selective and mechanistic pressures shaping their flight motor systems remain poorly understood. Here, we present a comprehensive comparative analysis of flight in Diptera (true flies), integrating morphology, wingbeat kinematics, and aerodynamics within a phylogenetic framework. We quantified morphology in 133 species spanning the Dipteran phylogenetic and size range, and for a subset of 46 species we combined high-speed stereoscopic videography with computational fluid dynamics (CFD) to characterize wingbeat kinematics and aerodynamic performance, respectively. Our results reveal that morphology is strongly structured by phylogeny, whereas wingbeat kinematics are broadly conserved across Diptera, reflecting dominant aerodynamic constraints. Two early-diverged lineages, Culicomorpha (mosquitoes and midges) and Tipulomorpha (crane flies), exhibit strikingly divergent kinematics and aerodynamics, suggesting lineage-specific selective pressures. Combining these data with our scaling analyses suggests that tiny Diptera are primarily constrained by aerodynamic force production, and maintain weight support through relatively larger wings and increased wingbeat frequencies. In contrast, as Diptera increase in size, hovering flight becomes progressively constrained by power availability, resulting in markedly elevated relative flight-muscle mass among the largest species. Mosquitoes and midges represent an extreme case, exhibiting a pronounced aerodynamic–acoustic trade-off with disproportionately large flight musculature and increased aerodynamic and acoustic power, consistent with selection favoring acoustic signaling during in-swarm mating. By integrating comparative morphology, kinematics, and aerodynamics across a major insect radiation, our study uncovers the interplay between physical scaling laws, aerodynamic constraints, and ecological pressures in shaping the evolution of animal flight. These findings provide a mechanistic framework for understanding how complex locomotor systems diversify under multiple selection pressures.

## Introduction

Insects are the most species-rich group of animals, comprising an estimated 5 million species – about 80% of all known animal species [1]. Their remarkable radiation may be partly attributed to their flight ability [2], which has led to a striking diversity of aerial lifestyles [3]. Despite the pivotal role that flight has played in the evolution and ecological importance of insects, our understanding of the evolutionary processes that generate the diversity of their flight kinematics, morphology, and associated aerodynamics remains limited.

This is in part because insect flight is remarkably complex and challenging to quantify [4], limiting the feasibility of large comparative evolutionary flight studies. While the flight of large vertebrate fliers can be addressed using classical aeronautical principles and often be inferred based on morphology [5–7], that of insects heavily relies on complex high-frequency flapping wing motions, resulting in unsteady aerodynamic mechanisms that fall outside the realm of conventional aerodynamic theory [4, 8, 9]. Precisely quantifying the diversity of wing morphology and wingbeat kinematics as well as the underlying unsteady aerodynamics is therefore crucial to understand the evolution of flight in insects.

Fundamentally, flying insects generate aerodynamic forces through the reciprocal motion of the wings swept at a high angle-of-attack, but the overall wingbeat kinematics may greatly vary among taxa. The extent to which the evolution of wingbeat kinematics is channeled by aerodynamic constraints, and how these constraints mediate flight divergence between species, is poorly understood. Although bounded by physical laws, variations in wingbeat kinematics among flying insects can be influenced by distinct evolutionary pressures. On one hand, neutral divergence along phylogenetic tree branches may lead to different flight kinematics between species. On the other hand, selective pressures acting on flight can strongly constrain the evolution of wingbeat kinematics, overcoming the pattern expected under neutral evolution.

Because flight is among the most energetically costly forms of locomotion [10], minimizing its energetic demands may represent one of the strongest selective pressures shaping the insect flight motor system. Cost-reducing selective pressures may be especially strong in highly aerial species, such as long-distance migrants [11]. Conversely, adaptation to some ecological niche may promote the evolution of specialized flight abilities, leading to wingbeat kinematics that diverge from those predicted by energetic optimization alone. For example, the evolution of aerial predatory habits in some groups (e.g., dragonflies or robber flies) may promote adapted flight kinematics and morphology enabling extremely agile flight [12, 13]. In contrast for prey species, natural selection may favor traits that enhance escape success, such as the rapid escape maneuverability of flies and mosquitoes [14, 15], or unpredictable flight behavior in mosquitoes and Lepidoptera [16, 17]. Sexual selection may also exert a powerful influence on the evolution of flight. In hoverflies for example, maintaining a highly stable stationary flight is crucial for guarding a mating territory [18, 19], which may drive the evolution of specialized hovering flight kinematics through male-male competition for females [19, 20]. Moreover, many insect species mate in swarms, and in some of the swarming species, males locate females by detecting the auditory flight tones produced by their beating wings [18, 21, 22]. It has been suggested that the high-frequency wingbeat produced by swarming mosquitoes and midges (Culicomorpha) results from sexual selection, as it maximizes audibility during mating interactions [21, 23, 24].

Flying insects span an impressive size range, from the tiniest 10 *µ*g fairy-fly to the largest 100 g Goliath beetle, covering seven orders of magnitude in body mass. Physical scaling laws strongly shape how the flight motor system functions across this range. One important effect is that, under geometric similarity, the ability of flapping wings to generate sufficient aerodynamic force for weight support decreases with decreasing body size. Tiny insects need to compensate for this through allometric adjustments in wing morphology or wingbeat kinematics [20]. As a consequence, the flight style that minimizes aerodynamic power differs across the size spectrum.

Aerodynamic scaling across sizes is expressed by the Reynolds number (Re), which quantifies the relative importance of inertial and viscous forces [25]. Flying insect wings operate at Reynolds numbers from tens to several thousands (Re ≈ 10–10^4^) [26]. This range places most species in the intermediate Re-regime, between a high-Re inertial regime where inertial forces dominate, and a low-Re laminar-viscous regime where viscous forces dominate [27]. These flow regimes strongly affect the aerodynamic drag (*D*) experienced by a flapping insect wing. In the inertial regime, drag is relatively low and approximately independent of Reynolds number (*D* ∝ Re^0^). In contrast, tiny insects operating near the laminar–viscous regime experience elevated drag because viscous forces dominate and the laminar boundary layer thickens. Under classical Blasius laminar boundary-layer theory, drag scales here inversely with Reynolds number as 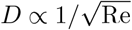 [27]. In the intermediate regime, drag therefore follows a scaling that lies between these two asymptotic limits. As a consequence, drag increases progressively as insect size decreases [27, 28].

This Re-dependent shift alters how drag, lift, and aerodynamic power scale with body size, and thus constrains the biomechanical strategies available to sustain powered flight. For example, tiny ptiliids beetles have evolved very lightweight bristled wings that yet still function effectively as solid surfaces at the low Reynolds numbers in which they operate [29]. In contrast, large flying insects such as *Morpho* butterflies, experience Reynolds numbers high enough to permit efficient gliding, and have evolved wing shapes adapted for this flight mode [30].

The diversity of insect flight kinematics likely results from the interplay between shared ancestry, physical scaling laws, aerodynamic constraints, and ecological pressures. Disentangling the relative role of these factors on the evolution of such a complex trait is a major challenge necessitating a coupled comparative evolutionary, biomechanical and aerodynamic perspective. The integration of these approaches has nevertheless rarely been attempted.

Contrasting with the huge number of flying insect species, flight has been comprehensively studied in only a handful of them, mostly with the aim to understand the aerodynamic mechanisms involved [14, 31–36]. Although providing valuable insights into flight mechanics [9], such studies generally consider single (model) species, or at most few species in comparison, and are thus of limited evolutionary relevance. In contrast, more extensive comparative evolutionary studies on insect flight mostly focused on morphology only, or simple flight kinematics parameters such as wingbeat frequency, and simplified aerodynamic modeling [3, 37–39]. This prevented us from fully appreciating the rich diversity in morphology, wingbeat kinematics and aerodynamics across taxa. The combination of detailed biomechanics studies and comparative phylogenetics is thus needed to understand the mechanistic and selective forces shaping the diversity of insect flight.

To address this knowledge gap, we performed a broad-scale comparative study of detailed morphology, wingbeat kinematics and aerodynamics, focusing on the order Diptera (true flies). Flies represent one of the largest radiation of flying animals [40]. With approximately 160,000 named species spread across *ca*. 180 families [41], and an estimated *>*1 million undescribed species, they represent approximately 15% of all animal species [1, 42]. Dipteran insects are found nearly everywhere on earth, inhabiting ecosystems ranging from the arctic tundras to arid deserts [43, 44]. They have been described as the ecologically most diverse animal order [45], as they comprise a wide array of trophic specializations (including but not limited to saprophages, predators, blood-feeders, herbivores, or pollinivores), as well as highly diverse sexual behaviors among species (including swarming, mate guarding, or male-male territorial contest) [46]. Moreover, the phylogenetic relationships are relatively well-resolved for most Dipteran taxa [41, 42, 47], enabling to investigate the macro-evolutionary processes shaping trait diversity across species (e.g., [48–50]). Finally, flight is central to many of Diptera’s essential behaviors for survival and reproduction, making this group an ideal model for addressing the evolutionary and biomechanical drivers of flight diversity.

Diptera are regarded as highly-specialized fliers, with a remarkably sophisticated sensory-motor system. Their flight is powered by asynchronous indirect flight muscles that drive rapid wingbeats, while tiny steering muscles at the wing base provide precise flight control and steering [51, 52]. Another key trait in Diptera is the presence of halteres, which are specialized sensory and flight control organs derived from the hindwings [48, 53]. These club-shaped structures confer a precise control over body rotations during flight. The combination of specialized sensory and motor systems make Dipteran insects among the most agile flying animals [52, 54].

Despite the high flight specialization of Diptera, marked differences in wingbeat kinematics exist among taxa. Most notably, the flight of mosquitoes (Culicidae) is characterized by low wingbeat amplitudes and high wingbeat frequencies [35, 55]. In contrast, other similarly-sized Diptera such as fruit flies (Drosophila) exhibit much larger wingbeat amplitude but lower frequencies [32]. Moreover, mosquitoes exhibiting slender body forms and high-aspect-ratio wings, while fruit flies possess stouter bodies and low-aspect-ratio wings. In addition, the relative mass of flight musculature varies significantly between these taxa, from approximately 30% of total body mass in Drosophila to 55 % in Culicidae species [56, 57], suggesting substantial variation in relative flight power availability. Mosquitoes and fruit flies thus represent two extremes in the diversity of Dipteran flight-motor systems. Beyond these two highly divergent species (among the earliest and most recently diverged taxa, respectively), the evolution of flight kinematic in most other Dipterans remain largely unexplored, particularly regarding the interplay between aerodynamic constraints, energetic flight efficiency, and other selective pressures.

In this study, we aim to uncover the evolutionary and biomechanical drivers of flight diversity in Diptera. For this, we quantified morphology, flight kinematics, and aerodynamic properties across a broad phylogenetic and size range of Diptera. Hereby, we use hovering flight as a standardized benchmark because it provides a consistent aerodynamic reference state across species.

Our investigation is structured around the following three objectives:

i. We ask whether the evolutionary diversification of Diptera flight was driven by large plasticity in wingbeat kinematics, or whether aerodynamic constraints channeled the evolution of flight kinematics. Such channeling would promote greater morphological than kinematic diversity across the order.
ii. We test how the physical requirement for powered flight is maintained across the vast size range of Diptera, evaluating whether this is achieved through allometric scaling of morphology, kinematics, or a combination of both. More specifically, we examine three key requirements for powered flight: generating sufficient aerodynamic force for weight support, minimizing aerodynamic power consumption, and ensuring that flight muscles can deliver the mechanical power needed for sustained flapping.
iii. We investigate whether lineage-specific selective pressures, such as selection for acoustic signaling in mosquitoes, promote the evolution of traits that depart from energy cost minimization. Such pressures may generate a trade-off between minimizing aerodynamic power and meeting conflicting demands imposed by other sexual or ecological selection.

By contrasting theoretical expectations from physiological and aerodynamic models and the observed flight diversity, our work highlights how phylogeny, sexual selection, and ecological pressures influenced the evolution of the Dipteran flight motor system. This comparative, phylogenetically informed, and size-aware approach enables us to evaluate lineage-specific traits against baselines defined by *similarly sized* Diptera. In doing so, it separates traits that are extreme in absolute terms from those that are unusually extreme for their size, and allows trait associations to be evaluated as broad comparative patterns instead of species-specific observations.

## Materials and methods

### Insect sampling

We quantified diversity in Diptera flight motor systems by sampling 133 species across 43 families spanning both Diptera’s phylogeny and body size range (Fig. 1A,B). Most species (n = 94) were collected near Wageningen University, the Netherlands (51°5901.0N, 5°3932.7E; altitude ca. 10 m) during the summers of 2021 and 2022. Additional sampling in the Amazonian rainforest of French Guiana (4°3409N, 52°1304W; ca. 300 m) in July 2021 yielded 36 species. Three mosquito species were obtained from laboratory rearing. The number of species per family was 3.2±3.7 (mean±standard deviation), with 1.9±1.2 individuals per species.

**Fig 1.**
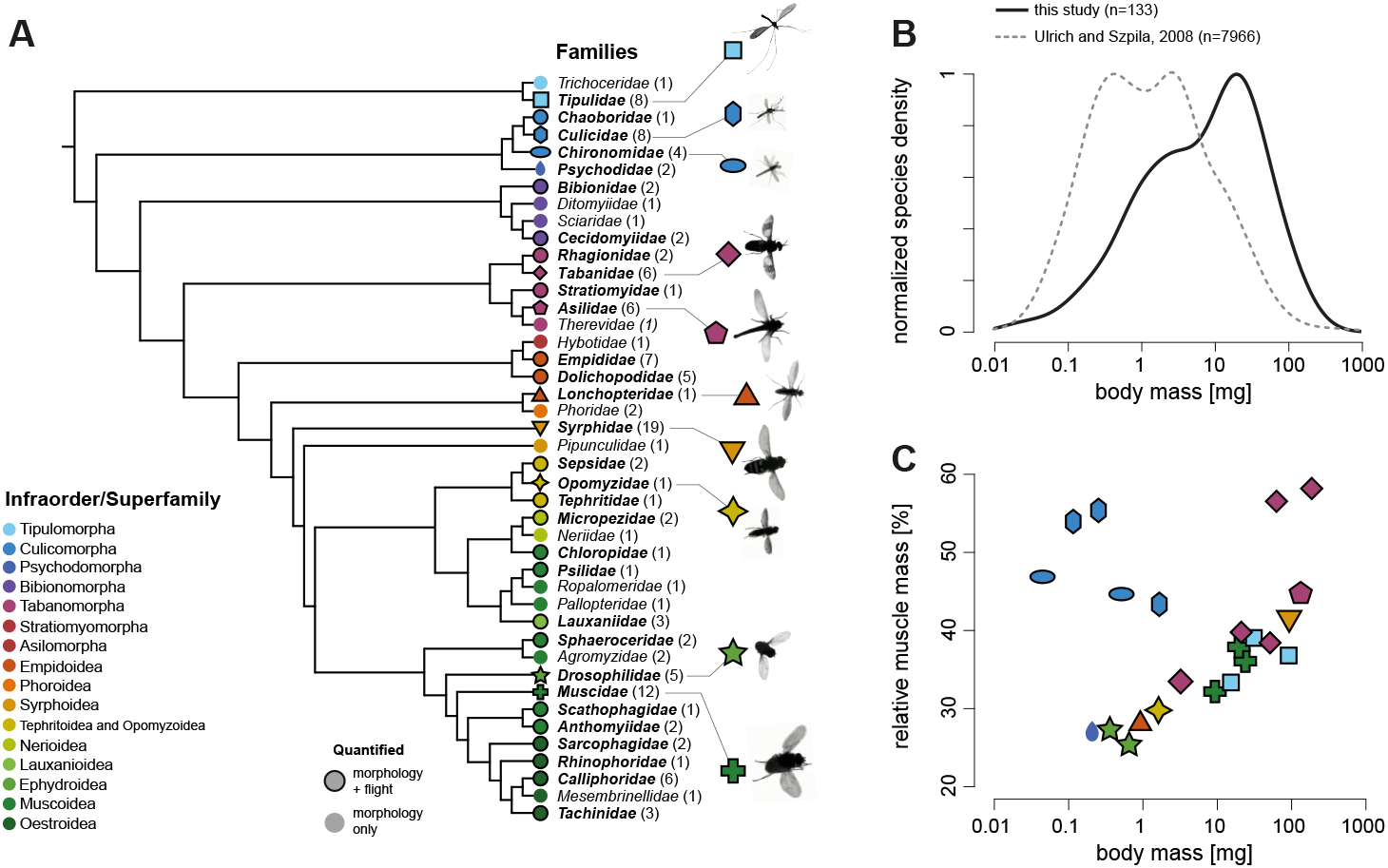
Phylogeny and morphology of the 133 studied Dipteran species. (**A**) The studied Diptera families shown with their phylogenetic relationships; the number of studied species for each family are given in parentheses. The families for which flight was quantified in at least one species are highlighted with black outlines. Phylogeny was obtained from Wiegmann *et al*. [41]. (**B**) Distribution of body mass in the sampled species (continuous black line) overlaid with a representative distribution of body mass across most Diptera (dashed grey line), estimated from a large meta-analysis covering nearly 8,000 species [59]. (**C**) Relative muscle mass *versus* body mass in a subset of species broadly spanning the phylogeny. The data points match non-circle symbols in (**A**). Species are colored by infraorder and/or superfamily to provide a clear overview of phylogenetic position. These groupings were chosen for visualization purposes and do not imply equivalent taxonomic rank.

Species were identified through DNA barcoding by submitting a leg from each individual to the Canadian Centre for DNA Barcoding, where standard automated BOLD protocols were used [58]. Sequences were matched to the BOLD reference library to establish species identities. We used the molecular phylogeny of Wiegmann *et al*. [41] and pruned it to the 133 sampled species (Fig. 1B).

Sample sizes per species were too low to test for sexual dimorphism. We therefore did not determine sex for most taxa. Culicomorpha were analyzed as males only because these taxa show strong sexual dimorphism in wingbeat kinematics and auditory behavior. Males form swarms, locate females phonotactically, and detect females through the interference (difference) tone generated by the beating wings during close-range interactions [21, 23, 24]. This male-specific behavior provides biological relevance for aero–acoustic comparisons but limits sex-specific inference.

Our sampling targeted broad size diversity but was slightly biased toward larger species, likely because larger individuals are more conspicuous in the field. Even so, the body-mass distribution in our dataset approximates that of Diptera as a whole, based on a meta-analysis of nearly 8000 species [59] (Fig. 1B).

For all species we quantified external body and wing morphology. For 24 species spanning 11 families we measured flight-muscle mass (Fig. 1C). For 46 species spanning 30 families we recorded hovering flight kinematics using stereoscopic high-speed videography and quantified aerodynamic forces and power using CFD simulations. For visualization, species were grouped by major lineages (infraorders or superfamilies), but all phylogenetic comparative analyses were conducted at the species level. Full sampling details are provided in Supplementary Methods S1.1.

### Quantifying body and wing morphology

We euthanized individuals by freezing at −20°C and measured fresh body mass and body-part masses on analytical balances appropriate to specimen size. For French Guiana specimens, fresh mass was estimated from dry mass using the relationship *m* = 4.1 *m*_dry_ − 0.96 (S1 Fig). We quantified flight-muscle mass for 24 species spanning a wide phylogenetic and size range by dissecting thoraxes (5.0±1.2 per species) and converting thorax mass to muscle mass as *m*_muscle_ = 0.9 *m*_thorax_.

Whole bodies and detached wings were photographed under a digital microscope, and body length and maximum thorax width were extracted from the calibrated images. Wing images were processed to obtain wing surface area *S*, wingspan *b*, and the second moment of area about the hinge *S*_2_ (Fig. 2A). From these we computed mean chord 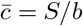, aspect ratio AR = *b*^2^*/S*, radius of gyration 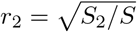, and its span-normalized form 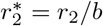.

**Fig 2.**
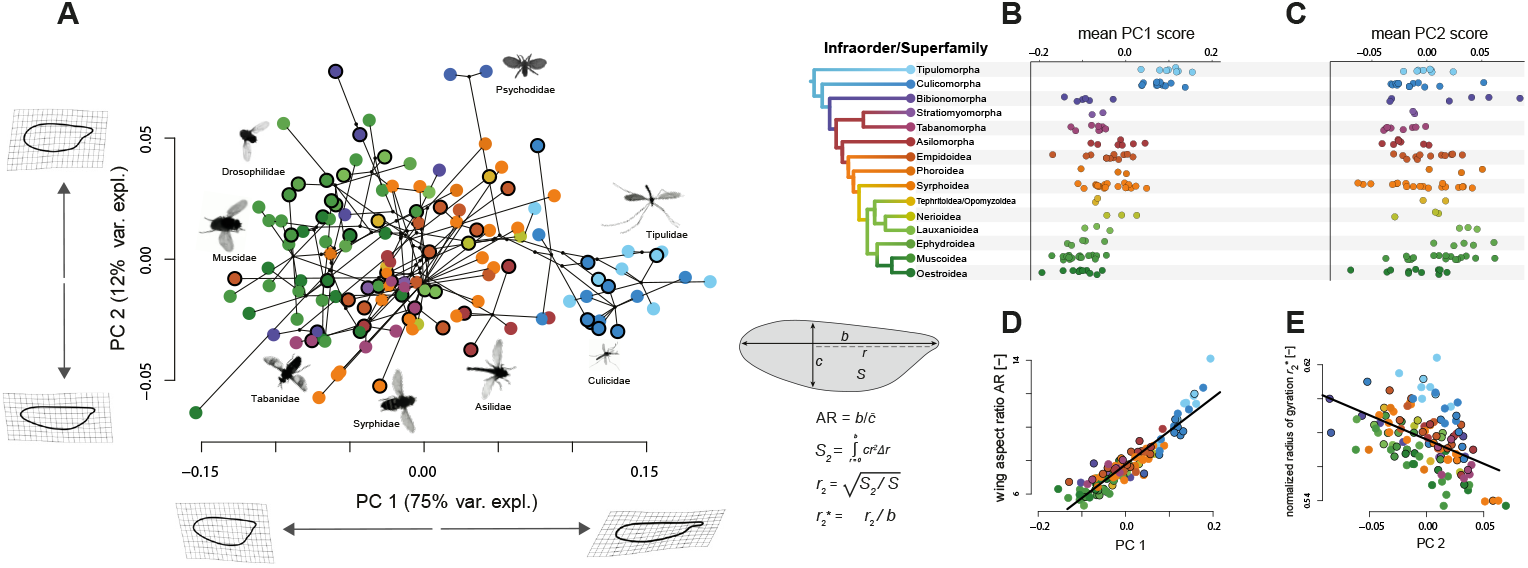
Wing shape variation across Diptera, quantified using a Principal Component Analysis (PCA). (**A**) The two first principal components (PC1 and PC2), together with the associated shape changes and percentage of the variation explained by the principal components. Shown wing shapes represent the extreme values associated with each PC axis. Each data point shows a species-average wing shape. Species for which flight was quantified are highlighted with black circles. (**B-C**) PC1 was strongly structured by phylogeny, while PC2 showed a weaker phylogenetic signal. (**D-E**) The two functional shape descriptors — aspect ratio (AR) and normalized second-radius-of-gyration 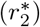 — were most strongly associated with PC1 and PC2, respectively.

To quantify wing-shape diversity, we used a landmark-based geometric morphometric approach. Each wing outline was represented by 300 sliding semi-landmarks and one fixed hinge landmark. After Procrustes superposition, we summarized shape variation across species using principal component analysis and visualized phylogenetic structure by overlaying the tree on the morphospace (Fig. 2A).

Full details of mass measurements, image processing, and geometric morphometric procedures are provided in Supplementary Methods S1.2.

### Flight experiments

We recorded hovering flight for 46 Diptera species spanning 31 families (Fig. 1A and Fig. 3). All individuals were collected near Wageningen University and tested in a custom octagonal Plexiglas arena (50 × 50 × 48 cm). Three synchronized high-speed cameras provided a top view and two oblique side views, with the focal volume centred in the arena and illuminated using infrared backlighting (Fig. 3A).

**Fig 3.**
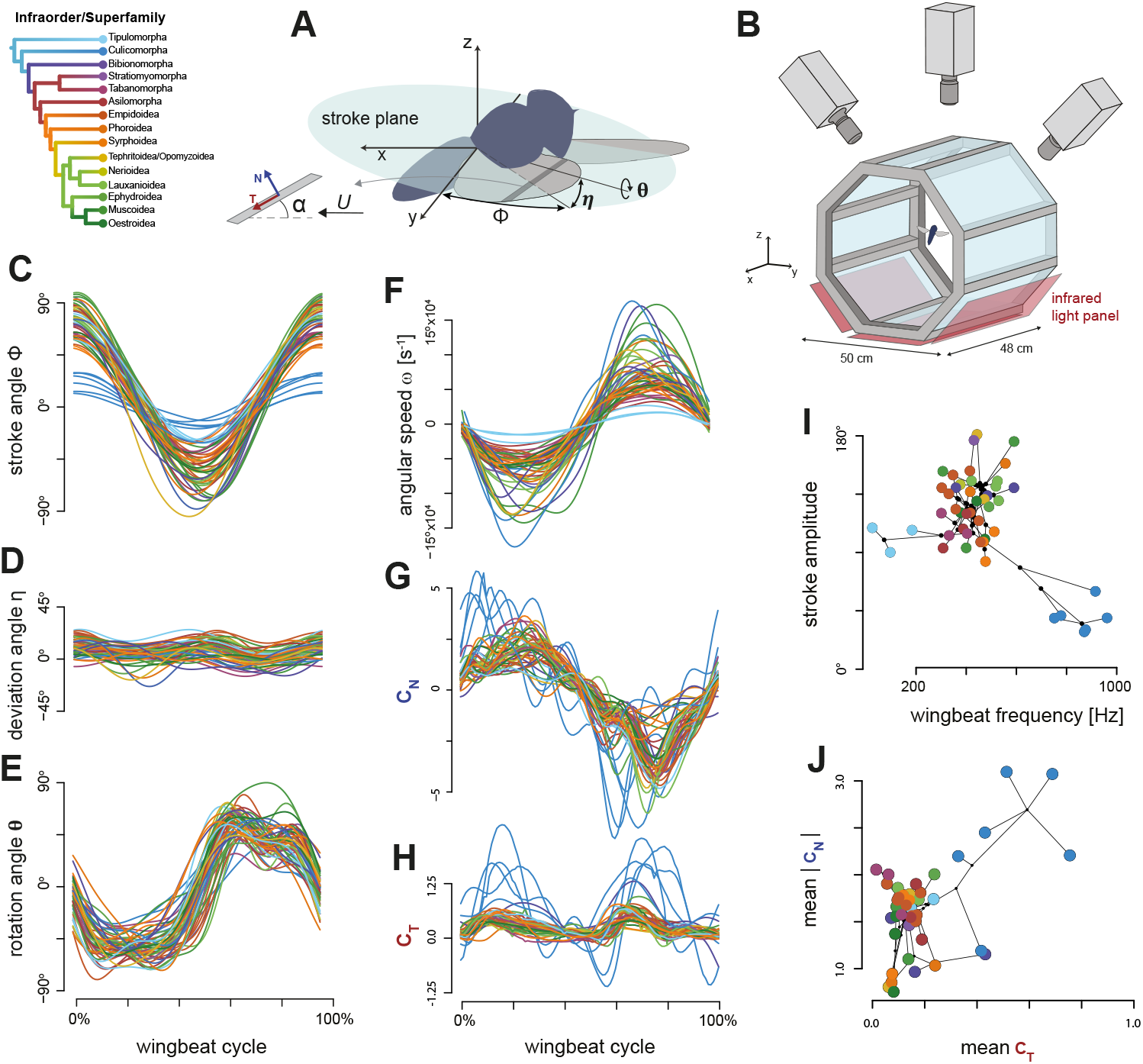
Wingbeat kinematics and aerodynamics of the 46 Diptera species studied in flight. (**A**) Definition of the wingbeat kinematics parameters: wing stroke angle within the stroke plane (*ϕ*), wing deviation angle out of the stroke plane (*η*), wing rotation angle along the spanwise axis (*θ*), air velocity relative to the wing (*U*), and wing angle-of-attack (*α*). Aerodynamic forces were estimated using CFD and are expressed as force coefficients normal and tangential to the wing surface (*N* and *T*). (**B**) Schematic of the flight experiment. Individuals were released into an octagonal flight arena and recorded with three synchronized high-speed cameras. From these recordings we reconstructed the three-dimensional body motion and wingbeat kinematics. Infrared backlighting provided high contrast between the flying insect and the background. (**C–H**) Temporal dynamics of wingbeat kinematics and aerodynamic force coefficients over a single wingbeat for all 46 species (S1 Video). Each trace shows species-averaged data and is color-coded by infraorder (legend top left). (**C–E**) Stroke, deviation, and rotation angles. (**F**) Angular speed. (**G**,**H**) Aerodynamic force coefficients normal and tangential to the wing surface. (**I**,**J**) Wingbeat-averaged kinematic and aerodynamic parameters for all species, color-coded by infraorder and connected by phylogeny. (**I**) Stroke amplitude versus wingbeat frequency. (**J**) Wingbeat-averaged absolute normal force coefficient versus the tangential force coefficient.

We used two optical configurations to accommodate the size range of Diptera. Larger species were filmed with lenses providing a focal zone of approximately 12 × 12 × 12 cm, and smaller species with a focal zone of approximately 6 × 6 × 6 cm. Frame rates were adjusted per species to achieve at least 20 frames per wingbeat; low-frequency fliers were recorded at ≥5,000 frames s^−1^, and high-frequency fliers at up to 13,500 frames s^−1^.

Before experiments, the stereoscopic system was calibrated using a T-wand moved throughout the focal zone, and DLT coefficients were computed from the tracked wand positions. During experiments several individuals were released simultaneously to increase the chance that one flew through the focal volume. When this occurred, recordings were saved using an end-trigger, and filming continued until at least three sequences were obtained for each species. Because direct assignment of flight measurements at the individual level were not possible, we here aimed at determining the average wingbeat kinematics per species instead of per individual. After experiments, all insects were euthanized and their morphology quantified as described above.

Full details on camera hardware, optical configurations, calibration procedures, and buffer-triggering are provided in Supplementary Methods S1.3.

### Quantifying body and wingbeat kinematics

We reconstructed 3D body pose and wing motion from stereoscopic videos using a validated MATLAB tracker adapted per species. Species-specific rigid wing models were derived from microscope images; wing deformations were not modeled. For each frame we obtained body position **X**(*t*) = [*x, y, z*] and body Euler angles (yaw *ψ*, pitch *β*, roll *φ*), and expressed wing motion in the stroke-plane frame using the Euler angles stroke *ϕ*, deviation *η*, and rotation *θ* (Fig. 3A,B).

To ensure temporal resolution across taxa, recordings provided on average 34.6 ± 7.7 frames per wingbeat; for low-frequency fliers some sequences were down-sampled. Wing-angle time series were fitted with fourth-order Fourier series, from which we computed wingbeat frequency *f*, stroke amplitude *A*_*ϕ*_ = *ϕ*_max_ − *ϕ*_min_, angle of attack *α*, and angular speed 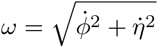.

Body-level kinematics per wingbeat included mean flight speed *U*, climb angle *γ*_climb_ = atan(*U*_ver_*/U*_hor_), and body pitch *β*_body_. We assessed proximity to hovering via the advance ratio 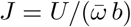, where 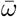 is wingbeat-averaged angular speed; *J <* 0.1 indicates hovering conditions.

Because flight measurements are variable, we quantified robustness of species-level differences using nested ANOVAs (sequences within species as a random effect) and reported inter-versus intraspecific coefficients of variation (Supplementary S2 Fig). Full tracking, optimization, and statistical details are provided in Supplementary Methods S1.4.

### Assessing the influence of phylogeny on morphological and flight traits

We tested whether evolutionary relatedness among Diptera influences variation in morphology and wingbeat kinematics by quantifying phylogenetic signal using Blomberg’s *K* for each parameter [60] and its multivariate extension [61]. Analyses were performed in *phytools* [62] using the Diptera phylogeny of Wiegmann *et al*. pruned to the species in our datasets (133 species for morphology; 46 for flight). To evaluate size trends, we fitted phylogenetically informed regressions (PGLS) on log_10_-transformed variables, estimating Pagel’s *λ* for each model. We report slopes, intercepts, confidence intervals, and the inferred phylogenetic signal.

This PGLS framework controls for shared ancestry and reduces leverage from clade-level outliers. Furthermore, by accounting for phylogenetic redundancy, it prevents closely related species from disproportionately influencing parameter estimates. Consequently, this allows statistically supported trends to be separated from patterns that are only suggested visually.

### Computational fluid dynamics simulations

We quantified aerodynamic forces and torques using full-DNS CFD simulations of a rigid, flat wing executing species-specific wingbeat kinematics. Each simulation used the measured wing outline and imposed the recorded wingbeat time series. The computational domain measured 10*b* × 10*b* × 10*b*, wing thickness was set to 0.025 *b*, and three wingbeat cycles were simulated. Because we focused on hovering flight, no mean inflow was imposed (*U*_∞_ = 0).

Simulations were performed with the open-source solver WABBIT [63, 64], which solves the incompressible Navier–Stokes equations using explicit finite differences with an artificial-compressibility formulation and dynamic wavelet refinement. Local grid resolution increased automatically where flow gradients required it. Starting from a coarse base grid, refinement proceeded up to seven levels, yielding an effective resolution of 307 points per wingspan. An eighth-level refinement test confirmed numerical convergence. Previous validation studies show that this solver produces quasi-exact solutions for rigid-wing flapping kinematics [63].

All simulations were run on distributed-memory clusters using up to 600 CPU cores. Full numerical details, refinement parameters, and validation tests are given in Supplementary Methods S1.5.

### Modeling the aerodynamics and aeroacoustics of Diptera flight

Generating sufficient lift while limiting energetic cost constrains the evolution of flapping-wing systems [25], and in several Diptera lineages sound production during swarming introduces additional selection pressures on wingbeat kinematics [65]. To interpret how morphology and kinematics shape aerodynamic force and power production and aeroacoustic output across species, we combined our full-DNS CFD simulations with a dimensional, mechanism-agnostic scaling framework. Below we summarize this framework; details are provided in Supplementary Methods S1.6.

#### Aerodynamic force production: CFD-validated scaling

We model the mean aerodynamic force during flapping flight using a minimal, dimensionally consistent representation that captures the key dependencies on morphology, kinematics, and aerodynamics. Following Buckingham–Pi analysis [66], the characteristic force scale for an oscillating wing is

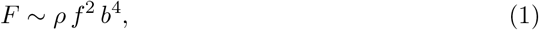

where *ρ* is air density, *f* wingbeat frequency, and *b* wingspan. Incorporating wing geometry and kinematic parameters yields the compact expression

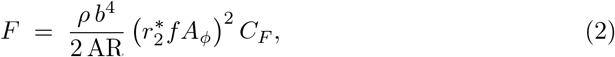

with aspect ratio AR = *b*^2^*/S*, span-normalized radius of gyration 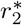, stroke amplitude *A*_*ϕ*_, and the wingbeat-average aerodynamic force coefficient *C*_*F*_. This formulation shows how wing size (*b*), wing shape (AR and 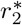), wingbeat kinematics (*f, A*_*ϕ*_), and aerodynamics (*C*_*F*_) combine to generate wingbeat-induced aerodynamic forces across species.

Depending on context, force vectors and coefficients were shown in one of two reference frames: (i) lift and drag, as perpendicular and parallel to the velocity vector, respectively; (ii) the wing-normal and tangential components (Fig. 3A). During steady hover the mean vertical force equals body weight, so each wing produces |*F*| = *mg/*2, giving the weight-normalized form

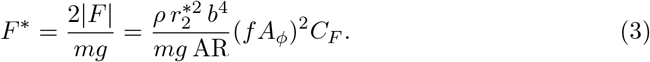

We validated this scaling against CFD-derived force coefficients and then used it to interpret how species of different sizes produce weight support during hovering flight.

#### Aerodynamic power for flight

Beyond generating sufficient lift, Diptera are expected to minimize the aerodynamic power required for flight. We quantified this power using full-DNS CFD, by computing instantaneous aerodynamic power per wing as

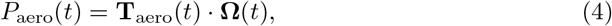

where **T**_aero_ and **Ω** are the aerodynamic torque and angular velocity vectors at the wing hinge. We then determined the wingbeat-averaged values, and normalized it with the mean aerodynamic force magnitude to obtain the weight-normalized wingbeat-average aerodynamic power as (in hover, |*F* | = *mg/*2)

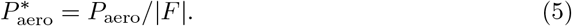

To interpret cross-species variation, we used a compact scaling surrogate derived by dimensional analysis. Assuming aerodynamic power is dominated by drag work, the weight-normalized aerodynamic power simplifies to

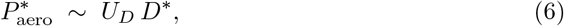

where *U*_*D*_ is the characteristic wing speed at the drag moment arm, and *D*\* = *C*_*D*_*/C*_*L*_ is the weight-normalized drag. Thus *U*_*D*_ captures the kinematic contribution to power, whereas *D*\* captures the aerodynamic contribution through lift–drag trade-offs.

We validated this surrogate against CFD-derived power for all species and used it only to interpret comparative trends. All aerodynamic forces and power values in this study come directly from CFD and therefore include unsteady, viscous, and Rankine–Froude-type induced-power effects. Weight-normalized aerodynamic power is reported as the flight-cost metric; it does not represent total mechanical power because inertial and elastic components are not included.

#### Acoustic power produced by flapping wings

Many Diptera species communicate acoustically during flight, especially in swarming taxa such as mosquitoes and midges [24, 67]. Aerodynamic sound arises from pressure fluctuations caused by wing motion. At the low Reynolds numbers of Diptera, where turbulence is limited, sound production is dominated by dipole sources generated by unsteady aerodynamic forces [68, 69]. The far-field acoustic pressure scales with the time derivative of the pressure-force component as

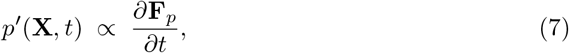

and integrating over a spherical surface yields acoustic power

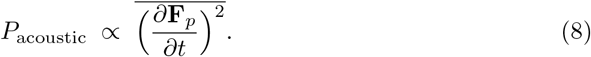

Dimensional reduction leads to a normalized acoustic-power metric,

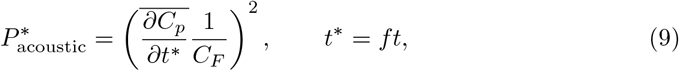

which separates pressure-coefficient fluctuations (the source of sound) from the aerodynamic force coefficient required for weight support. This metric therefore quantifies sound production per unit of aerodynamic force generation during hovering flight.

### Estimating available flight power from muscle morphology

We compared the previously estimated required aerodynamic power for flight 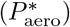 with the power available in the flight musculature 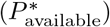. Weight-normalized available power was estimated as

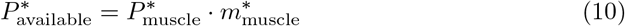

where 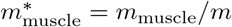 is the relative flight muscle mass, and 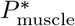 is the wingbeat cycle-averaged power output per kilogram of muscle. We used a value of 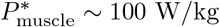 W/kg, consistent with previous studies reporting a relatively narrow range of 80–120 W/kg across insect taxa and body sizes [56, 70, 71]. As described above, we estimated the flight muscle masses based on thorax mass [56]. We then tested whether interspecific differences in power requirements were matched by corresponding variation in available power using a PGLS regression.

### Adaptations in morphology, wingbeat kinematics and aerodynamics across the Diptera phylogeny

We analyzed how morphological, kinematic, and aerodynamic traits vary across Diptera and how these traits contribute to weight support, aerodynamic cost, and wingbeat-induced sound production. We interpreted these patterns using the CFD-validated aerodynamic and acoustic scaling framework (Eqs. 3, 5, 8, 9; full derivations in Supplementary Methods S1.6–S1.9).

#### Scaling of aerodynamic force production with body mass

Weight support in hover requires that the mean aerodynamic force scales linearly with body mass (*F* ∝ *m*). Under geometric, kinematic, and dynamic similarity this requirement is not met: similarity predicts *F* ∝ *m*^4*/*3^ (Eq. 1; Supplementary Methods S1.8; [20]). Smaller species must therefore compensate through morphology, kinematics, or both. Using the aerodynamic force model, we defined the theoretical scaling expected if only a single parameter were adjusted to maintain weight support (*F* ∝ *m*). These scenarios predict, for example, *b*~*m*^1*/*4^, 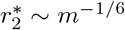, AR~*m*^1*/*3^, *f*~*m*^−1*/*6^, *A*_*ϕ*_~*m*^−1*/*6^, or *C*_*F*_~*m*^−1*/*3^ (see Table 1 for the full list; Supplementary Methods S1.8 for the derivations).

**Table 1.**
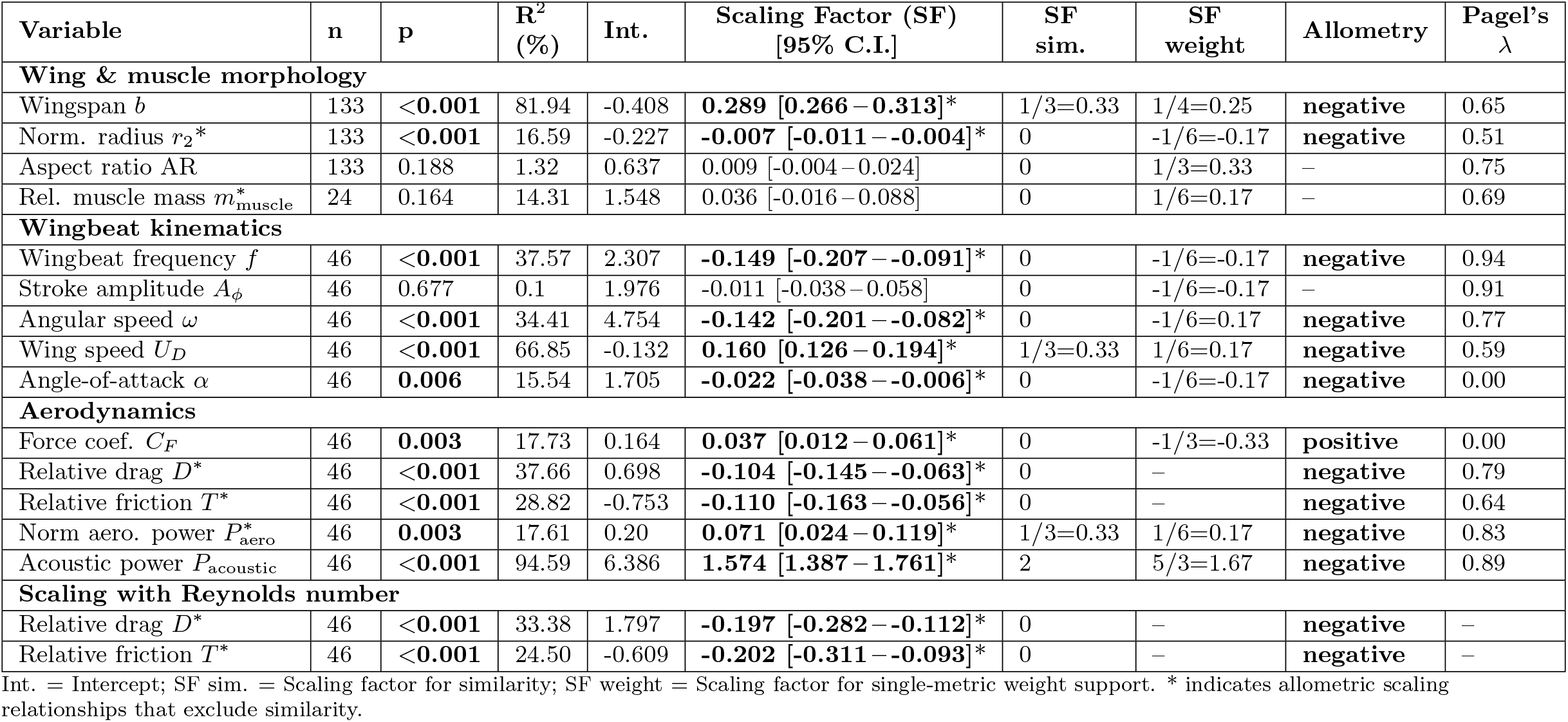
Results of phylogenetic general least square (PGLS) regressions of wing morphology, wingbeat kinematics and aerodynamic parameters. All regressions were performed using log10-transformed values, and relative to body mass, except for the bottom section with heading *Scaling with Reynolds number*. Stars indicate 95% confidence intervals that exclude the scaling factor for similarity, suggesting allometry.

| Variable | n | p | R<sup>2</sup> (%) | Int. | Scaling Factor (SF)<br>[95% C.I.] | SF<br>sim. | SF<br>weight | Allometry | Pagel's<br>$\lambda$ |
| --- | --- | --- | --- | --- | --- | --- | --- | --- | --- |
| **Wing & muscle morphology** |  |  |  |  |  |  |  |  |  |
| Wingspan $b$ | 133 | <0.001 | 81.94 | -0.408 | **0.289** [0.266 – 0.313]* | 1/3=0.33 | 1/4=0.25 | negative | 0.65 |
| Norm. radius $r_2^*$ | 133 | <0.001 | 16.59 | -0.227 | **-0.007** [-0.011 – -0.004]* | 0 | -1/6=-0.17 | negative | 0.51 |
| Aspect ratio AR | 133 | 0.188 | 1.32 | 0.637 | 0.009 [-0.004 – 0.024] | 0 | 1/3=0.33 | – | 0.75 |
| Rel. muscle mass $m_{\text{muscle}}^*$ | 24 | 0.164 | 14.31 | 1.548 | 0.036 [-0.016 – 0.088] | 0 | 1/6=0.17 | – | 0.69 |
| **Wingbeat kinematics** |  |  |  |  |  |  |  |  |  |
| Wingbeat frequency $f$ | 46 | <0.001 | 37.57 | 2.307 | **-0.149** [-0.207 – -0.091]* | 0 | -1/6=-0.17 | negative | 0.94 |
| Stroke amplitude $A_\phi$ | 46 | 0.677 | 0.1 | 1.976 | -0.011 [-0.038 – 0.058] | 0 | -1/6=-0.17 | – | 0.91 |
| Angular speed $\omega$ | 46 | <0.001 | 34.41 | 4.754 | **-0.142** [-0.201 – -0.082]* | 0 | -1/6=0.17 | negative | 0.77 |
| Wing speed $U_D$ | 46 | <0.001 | 66.85 | -0.132 | **0.160** [0.126 – 0.194]* | 1/3=0.33 | 1/6=0.17 | negative | 0.59 |
| Angle-of-attack $\alpha$ | 46 | **0.006** | 15.54 | 1.705 | **-0.022** [-0.038 – -0.006]* | 0 | -1/6=-0.17 | negative | 0.00 |
| **Aerodynamics** |  |  |  |  |  |  |  |  |  |
| Force coef. $C_F$ | 46 | **0.003** | 17.73 | 0.164 | **0.037** [0.012 – 0.061]* | 0 | -1/3=-0.33 | positive | 0.00 |
| Relative drag $D^*$ | 46 | <0.001 | 37.66 | 0.698 | **-0.104** [-0.145 – -0.063]* | 0 | – | negative | 0.79 |
| Relative friction $T^*$ | 46 | <0.001 | 28.82 | -0.753 | **-0.110** [-0.163 – -0.056]* | 0 | – | negative | 0.64 |
| Norm aero. power $P_{\text{aero}}^*$ | 46 | **0.003** | 17.61 | 0.20 | **0.071** [0.024 – 0.119]* | 1/3=0.33 | 1/6=0.17 | negative | 0.83 |
| Acoustic power $P_{\text{acoustic}}$ | 46 | <0.001 | 94.59 | 6.386 | **1.574** [1.387 – 1.761]* | 2 | 5/3=1.67 | negative | 0.89 |
| **Scaling with Reynolds number** |  |  |  |  |  |  |  |  |  |
| Relative drag $D^*$ | 46 | <0.001 | 33.38 | 1.797 | **-0.197** [-0.282 – -0.112]* | 0 | – | negative | – |
| Relative friction $T^*$ | 46 | <0.001 | 24.50 | -0.609 | **-0.202** [-0.311 – -0.093]* | 0 | – | negative | – |

Int. = Intercept; SF sim. = Scaling factor for similarity; SF weight = Scaling factor for single-metric weight support. \* indicates allometric scaling relationships that exclude similarity.

To evaluate how real Diptera achieve weight support, we fitted PGLS regressions for each morphological, kinematic, and aerodynamic parameter using log_10_-transformed values and the Diptera phylogeny. We then tested for each parameter whether the empirical PGLS scaling deviated from its similarity expectation and from its theoretical weight-support scaling. To do this, we examined whether the 95% confidence interval of the PGLS slope excluded the corresponding similarity and weight-support slopes. Parameters whose intervals did not overlap the similarity expectation were classified as allometric (Table 1).

Finally, we quantified the relative contributions of each trait to maintaining weight support across sizes by calculating its relative allometric scaling for weight support as [20]

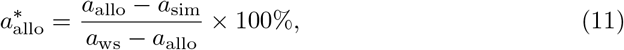

where *a*_allo_ is the observed allometric slope (PGLS), and *a*_sim_ and *a*_ws_ are the corresponding expectations under similarity and under single-trait weight support, respectively (see Supplementary Methods S1.9 for details).

#### Assessing the role of cost minimization in Diptera flight

We next tested whether Diptera minimize the aerodynamic power required for hover. Weight-normalized aerodynamic power was computed directly from CFD 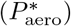 and interpreted using the scaling surrogate 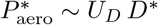 (Eqs. 4–6). Under weight-support constraints we expect *U*_*D*_ ~ *m*^1*/*6^, and under geometric and kinematic similarity *U*_*D*_ ~ *m*^1*/*3^. Assuming dynamic similarity (*D*\* ~ *m*^0^), this yields 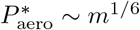. For each CFD-studied species we computed 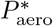, *U*_*D*_, and *D*\* and tested their mass scaling via PGLS, and compared these with the theoretical expectations using the confidence-interval approach described above.

Because Diptera operate in an intermediate Reynolds number regime (Re ~ 10^1^–10^4^), we also tested how *D*\* scales with Re. Classical fluid-mechanical expectations [27] and systematic insect-flight work [28] show that in this regime the drag coefficient transitions from Reynolds-number independence at the higher inertial Reynolds number regime (*D*\* ∝ Re^0^) to an inverse dependence at the lower laminar–viscous Re-regime 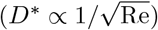 [27].

We tested for this effect in Diptera using a series of PGLS fits, following the approach described above. Hereby, we first tested how relative drag scales with Reynolds number. To isolate viscous contributions, we controlled for angle-of-attack by relating *D*\* to the wingbeat-average effective *α*, and then examining residual variation attributable to viscosity. We further quantified the viscous component using the relative tangential force on the wing *T* * = *C*_*T*_ */C*_*N*_, and its scaling with Re.

#### Scaling of aero-acoustic power with body mass and aerodynamic efficiency

To examine whether Diptera that use acoustic communication generate disproportionately high sound, and whether this incurs aerodynamic cost, we computed aero-acoustic power from CFD and analyzed its mass scaling. Theoretical expectations give *P*_acoustic_ ~ *m*^2^*f* ^2^ (Eq. S10); if frequency alone compensates for weight support (*f* ~ *m*^−1*/*6^), then *P*_acoustic_ ~ *m*^5*/*3^. We tested whether empirical PGLS slopes differed from this prediction using the confidence-interval approach described above.

To test for an aerodynamic trade-off, we computed the normalized acoustic-power metric 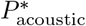 (Eq. 9) and assessed its relationship with the aerodynamic cost *D*\*. We then compared patterns between Diptera that rely on acoustic mate-location and those that do not. Acoustic derivations and normalization details appear in Supplementary Methods S1.6.

## Results

### Morphological and kinematic diversity in Diptera

To evaluate our three primary objectives, (i) the effect of aerodynamic constraints on kinematic diversity, (ii) the scaling of morphology, kinematics and aerodynamics with size, and (iii) the trade-off between flight cost minimization and acoustic signaling, we first quantified the range of phenotypic variation in Diptera.

We investigated morphological diversity across 133 Diptera species and flight kinematic diversity in a subset of 46 species (Fig 1, S1 Video). For this, we quantified the body and wing morphology of 220 individual insects (1.9±1.2 individuals per species), and the body and wingbeat kinematics throughout 276 single-wing wingbeats in 46 Diptera species (six single-wing wingbeats per species). The species studied broadly covered the phylogenetic tree (Fig 1A), and a large range of body masses, from 0.02 mg (*Micromya lucorum*, Cecidomyiidae) to 227 mg (*Tabanus sp*., Tabanidae) (Fig 1B).

#### Morphological diversity in Diptera

Our geometric morphometrics analysis revealed that wing elongation represents the primary axis of variation in the wing shape morphospace, accounting for 75% of the total shape variation (PC1 Fig 2A). Wing elongation differentiates the major dipteran groups well: early-diverging lineages exhibit elongated wings (positive PC1 values), whereas more recently diverged groups tend to have compact wings (negative PC1 values). Accordingly, variation in wing aspect ratio was strongly structured by phylogeny (Fig 2B,D), and exhibited the strongest phylogenetic signal among all morphological parameters (K = 0.69; S1 Table). PC2 accounted for 12% of the total shape variation, and primarily captured shape changes corresponding to variation in the normalized radius of gyration, 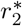 (Fig 2C,E). Higher PC2 values indicate greater distal distribution of wing area, while lower values reflect area concentrated near the wing base. This axis of shape variation scaled allometric with body mass (PGLS of PC2 against body mass: R^2^=0.16; t=-4.75, p *<*0.001) (Fig 2E). Although less clearly associated with phylogeny, 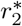 still exhibited a significant phylogenetic signal (Fig 2C; K = 0.24; S1 Table).

Interestingly, variation in body shape paralleled that of wing shape, as the correlation between wing and body aspect ratio was significant and positive (r=0.69; t=11.12, p *<*0.001), with early-diverged taxa tending toward elongated high-aspect ratio body and wing forms. Shape characteristics thus mainly differentiated Dipteran taxa, ranging from slender morphologies in early-diverged lineages to stouter bodied in recently diverged ones. In contrast, body size was markedly more variable within taxa and showed no significant phylogenetic signal (S1 Table). Here, references to “early-diverged” lineages pertain to extant taxa and should not be interpreted as statements about the phenotype of ancestral Diptera.

#### Diversity in hovering flight kinematics and aerodynamics among Diptera

The quantification of hovering flight kinematics effectively captured inter-specific variation, with differences between species mostly larger than individual variability across flight sequence within the same species (S2 Fig and S2 Table).

Overall, the wingbeat kinematics of the 46 Dipteran species were remarkably similar across the phylogeny, with the exception of the two early-diverged taxa: Culicomorpha (mosquitoes and midges) and Tipulomorpha (crane flies) (Fig 3, S1 Video). Amoung most Diptera taxa, wingbeat frequency is conserved within relatively narrow range of *f* = 236 [201–270] Hz (mean [95% confidence interval]; Fig 3I). In contrast, Culicomorpha and Tipulomorpha showed striking divergence, exhibiting wingbeat frequencies 300% and 30% of the median Diptera, respectively (Culicomorpha: *f* = 714 [536 – 924] Hz; Tipulomorpha: *f* = 70 [12 – 118] Hz; Fig 3C).

Focusing on the temporal dynamics of their wingbeat kinematics show similar trends (Fig 3C–E, S1 Video). In all species, the wingbeat involved a forward and backward wing movement following a sinusoidal pattern within the stroke plane (Fig 3C), with a relatively narrow range in stroke amplitude (*A*_*ϕ*_ = 117° [110°–124°]). Culicomorpha are here a clear outlier, with exceptionally low amplitudes (*A*_*ϕ*_ = 37° [32°– 42°]). Movements outside the stroke plane, reflected by the deviation angle, were for all Dipteran species minimal (Fig 3E). As such, the oscillatory angular speed of the wingbeats was primarily caused by the wing stroke movement, with peak of angular speed *ω* = 68° [62° – 74°] × 10^3^ s^−1^ (Fig 3F). Here, Tipulomorpha were the outlier, with exceptionally low wing stroke speeds (*ω* = 21°[14°– 27°] × 10^3^ s^−1^), caused by their low wingbeat frequencies. For most wingbeats, the wing maintained a relatively constant rotation angle during the mid forward and backward strokes, often exhibiting a ‘double peak’ pattern (Fig 3E). At the end of both the forward and backward strokes, the wings quickly supinated and pronated, respectively. This causes the wing to flip upside down at stroke reversal, allowing it to operate again at a positive constant angle-of-attack *α*, required for producing an upward-directed aerodynamic force (Fig 3G).

The conserved wingbeat pattern among most Diptera resulted in similarly conserved aerodynamic force production, as quantified by the force coefficients normal and tangential to the wing surface, derived from CFD (Fig 3G, H and J). Hereby, the Culicomorpha species are again strikingly different from the norm.

For the non-Culicomorpha Diptera, the temporal dynamics of the normal force coefficient (*C*_*N*_ (*t*)) shows two peaks (Fig 3G): a positive peak around mid forward wingstroke, and a negative peak on the supinated wing at mid backstroke. This results in upward-directed force production during both strokes, with a wingbeat-average absolute normal force coefficients of |*C*_*N*_ | = 1.54 [1.43 – 1.65] (Fig 3J). The temporal dynamics of the tangential force coefficient (*C*_*T*_ (*t*)) is also conserved among the non-Culicomorpha species, generating a tangential force peak at each mid wingstroke (Fig 3H), resulting in wingbeat-average tangential force coefficients of *C*_*T*_ = 0.13 [0.11 – 0.15].

In contrast, Culicomorpha species show distinct temporal dynamics in normal and tangential force production compared with the other taxa (Fig. 3G,H). Their force coefficients reach higher peak values, the normal force peaks earlier in the wingbeat cycle, and the tangential forces show negative thrust-producing peaks near stroke reversal. These patterns are consistent with earlier analyses of mosquito flight aerodynamics [35]. The elevated peak forces also produce significantly higher wingbeat-averaged force coefficients (Fig. 3J). In Culicomorpha, the wingbeat-averaged normal and tangential force coefficients are 54% and 300% higher than in the other taxa, respectively (|*C*_*N*_| = 2.37 [1.63 – 3.11] and *C*_*T*_ = 0.52 [0.34 – 0.69]).

Regarding general flight kinematics, the mean flight speed was *U*_∞_ = 0.76 [0.71 – 0.81] m s^−1^, while the mean body pitch and climb angle were *β*_body_ = 31° [28°– 34°]) and *γ* = 20° [16°– 25°], respectively. These two angles showed greater variability between individuals than between species, obscuring possible species-specific differences. Similarly, the stroke plane *β*_stroke-plane_ = −15° [−19° – − 11°] showed higher variability within than between species (see S2 Fig and S2 Table for inter-vs intra-specific variation in all flight parameters). The mean advance ratio was close to the generally accepted threshold of 0.1 (*J* = 0.13 [0.12 – 0.14]), showing that the studied Diptera were operating relatively close to a hovering flight mode.

Wingbeat kinematics showed significant phylogenetic signal for all parameters (*K*-multivariate = 0.01; p = 0.29), excepted for the angle-of-attack that was not influenced by phylogeny (see S1 Fig for parameter-specific values), suggesting strong aerodynamic constraints limiting its variation.

### Allometric adaptations in morphology and wingbeat kinematics for maintaining weight support across sizes

To address our second objective, the scaling of morphology and kinematics (ii), we evaluated how the fundamental physical requirement for weight support (2|*F* | = *mg*) is maintained across the vast size range of Diptera through allometric adaptations in wing morphology and wingbeat kinematics (Fig 4). This revealed that, among the morphological parameters that primarily affect aerodynamic force production (Eq. 3), wingspan (*b*) and normalized radius of gyration 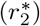 show negative allometric scaling with body mass (Fig 4A and B, respectively); in contrast, aspect ratio (AR) did not scale with size (Fig 4C). Thus, tiny Diptera species have relatively larger wingspans than larger species, and have more spatulated wings (high 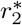). Both mechanisms allow tiny fliers to (partly) compensate for the reduction in aerodynamic force production under geometric similarity. While aspect ratio did not scale with size, it varied primarily with phylogeny (Fig 2), with the early-diverged taxa Culicomorpha and Tipulomorpha having exceptionally high aspect ratio wings (Fig 4C; among all Diptera: AR=3.86 [3.72 – 4.00]; among Culicomorpha and Tipulomorpha: AR=5.23 [4.93 – 5.54]).

**Fig 4.**
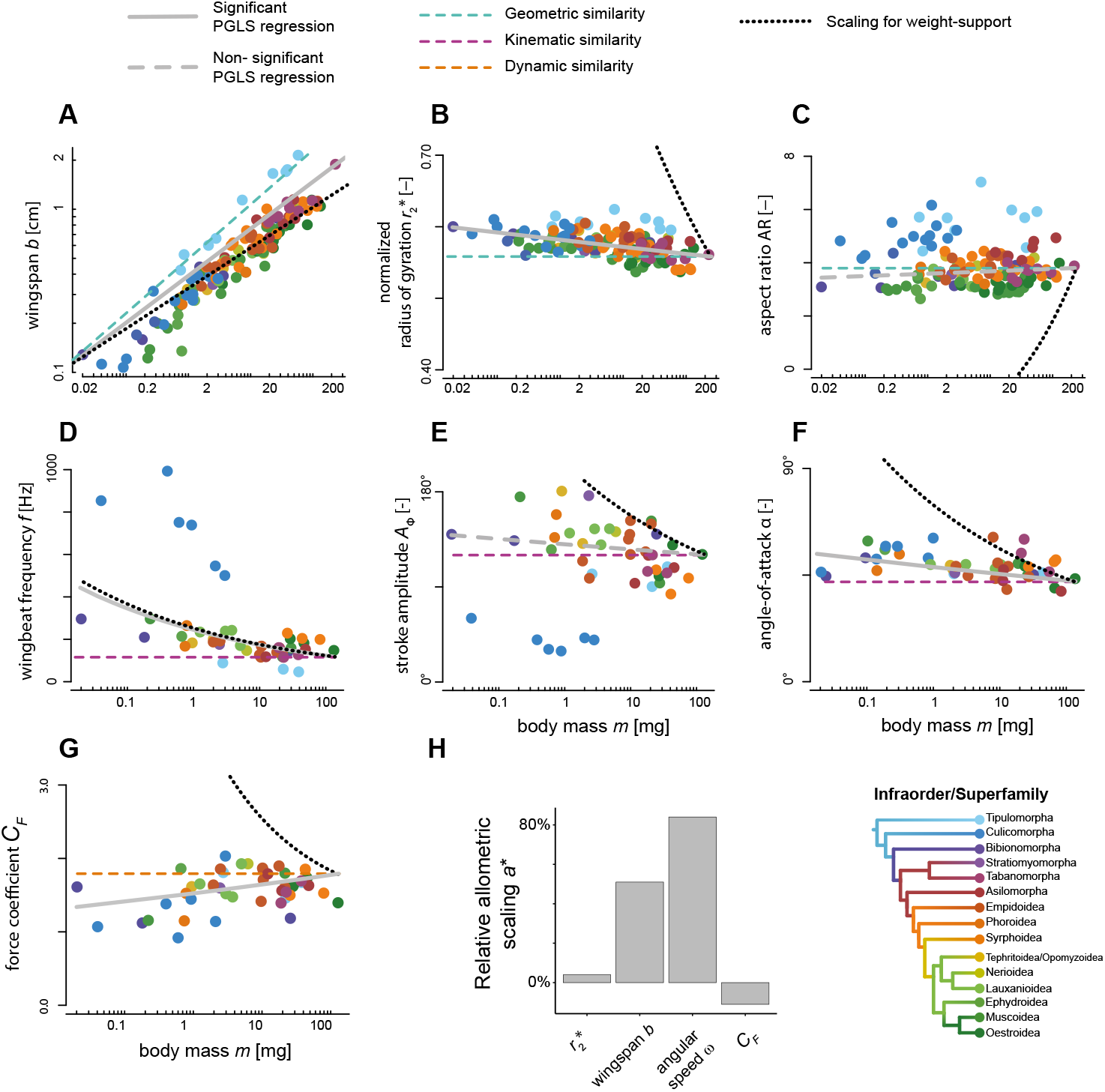
Scaling of morphology, wingbeat kinematic and aerodynamics with body mass. (**A-G**) Relationship between morphological, kinematics and aerodynamics parameters from the aerodynamic force model (Eq. 3) and body mass. Data points show species-average results, and are color-coded by infraorder (see bottom right). The best fit from the PGLS regression on the data is shown as a gray line; the trend line is solid when the parameter varies significantly with body mass, or dashed when the relationship is not significant. Expected slopes under geometric, kinematic, and dynamic similarity are shown as blue, purple, and orange dashed lines, respectively. The expected scaling for maintaining weight support (assuming all other parameter scale under similarity) is shown as a black dotted line. All plots are displayed on a linear–log scale, except for wingspan, which is shown on a log–log scale. (**H**) Relative contributions of the different parameters to maintaining weight support across sizes, as expressed by the relative allometric scaling parameter *a*\*.

Among the wingbeat kinematics parameters that affect aerodynamic force production (Eq. 3), both wingbeat frequency and angle-of-attack exhibited negative allometric scaling with body mass (Fig 4D and F, respectively); wingstroke amplitude did not scale with size (Fig 4E). Thus, tiny Dipteran fliers beat their wings at higher frequencies than larger species, and do so at higher wingbeat-average angles-of-attack; in contrast, they do not significantly increase their wingstroke amplitude. The increase in wingbeat frequency allow tiny fliers to further compensate for the reduction in aerodynamic force production for scaling under geometric similarity (Fig 4D). In contrast, the increase in angle-of-attack for tiny Diptera seems to cause a corresponding reduction in the wingbeat-average force coefficient, which show a positive allometric scaling with size (Fig 4G). This is expected because the angles-of-attack in tiny fliers are higher than 45°, where thus an increase in angle-of-attack leads to a reduction in the upward-directed force [72].

These combined results suggest that allometric adaptations in both morphology and wingbeat kinematics allow Diptera of different sizes to maintain weight support during hovering flight. We estimated the relative contribution of allometric adaptations in the different morphology and kinematics parameters to maintain weight support across sizes, by calculating the relative allometric scaling factor *a*\* (Fig 4H). This showed that allometric adaptations in wingbeat kinematics, as summarized by the angular wing speed, contributed the most to maintaining weight support across sizes (84%), which is primarily achieved via modulations in wingbeat frequency (88%). Allometric adaptations in wing morphology consisted primarily of an increase in wingspan with reducing size (contributing 51%), and lesser so by allometric adaptations in the normalized radius of gyration (4%). The scaling in the wingbeat-average force coefficient had a detrimental negative effect (−11%). The sum of all significant relative allometric scaling factors adds to 128%, and thus these metrics should be used in a relative manner, not in absolute sense.

### Scaling of aerodynamic power with size and phylogeny

In addition to maintaining weight support, flying Diptera are expected to minimize the power required for flight. We combined CFD with physical scaling law models (Eq. 6) to assess how aerodynamic power in hovering flight scales with size, phylogeny, kinematics and aerodynamics (Fig 5).

**Fig 5.**
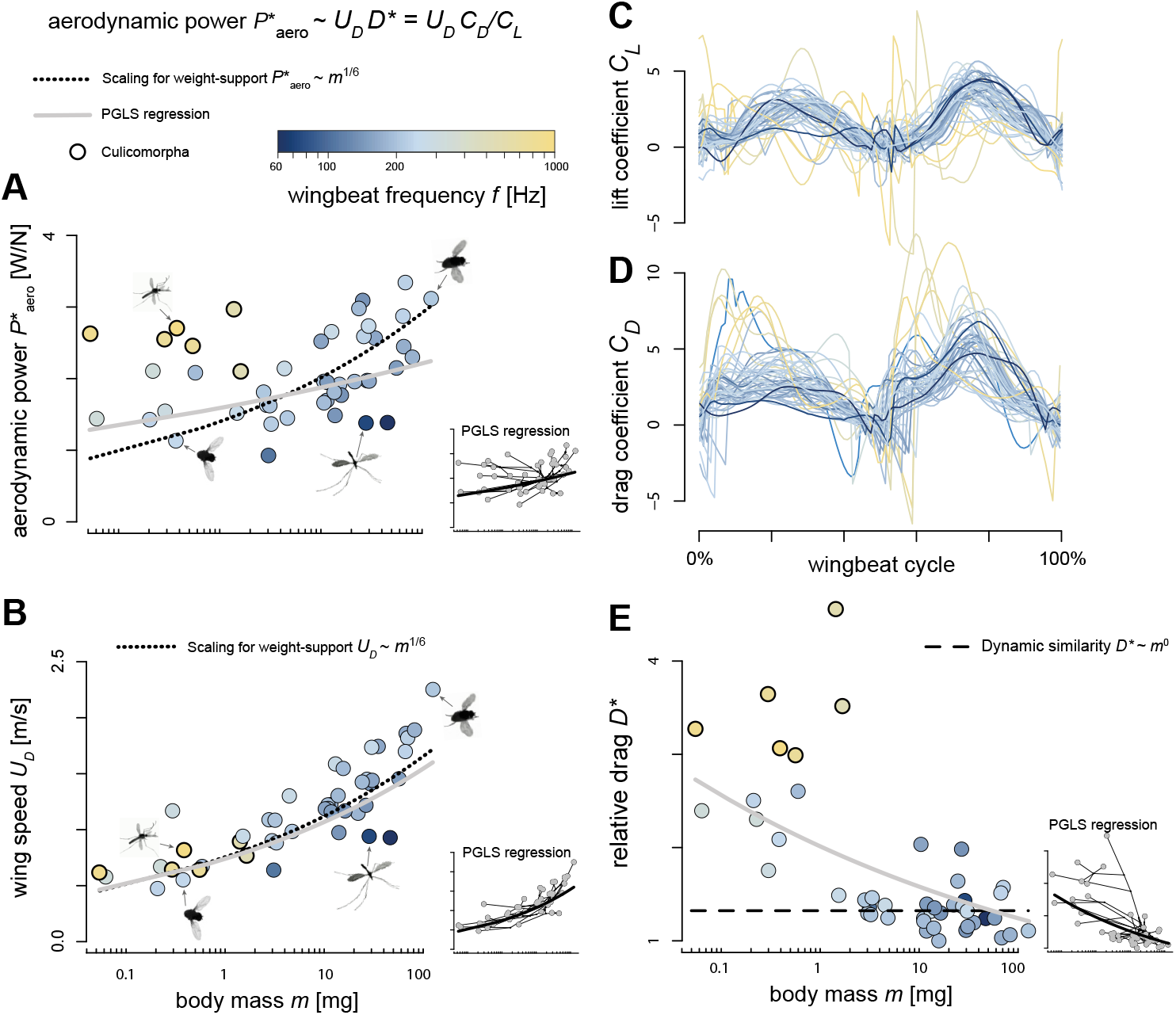
Scaling of aerodynamic power with body mass and temporal dynamics of lift and drag. Data points show species-average values and are color-coded by wingbeat frequency (top). Culicomorpha species with elevated wingbeat frequencies and swarm-based mating are highlighted in bold. Trend lines show PGLS regression fits (gray lines), scaling for weight-support (dotted lines), and for dynamic similarity (dashed). (**A**,**B**,**E**) Scaling of CFD-derived normalized aerodynamic power with body mass (**A**), and scaling of its two sub-components based on 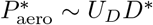, being wing speed (**B**) and relative drag (**E)**. (**A**) The significant PGLS-based scaling shows that aerodynamic power increases with body mass. Species with higher wingbeat frequencies show for their size elevated power requirements. (**B**) Wing speed increases with body mass, following the size dependence expected under weight-support constraints (*U*_*D*_ ~ *m*^1*/*6^). (**E**) Relative drag (*D*\* = *C*_*D*_*/C*_*L*_) increases with reducing size, particularly for small Diptera; relative drag is exceptionally high in high-frequency fliers. (**C**,**D**) Temporal dynamics of lift and drag coefficients for all species. Because *D*\* = *C*_*D*_*/C*_*L*_, these traces show that the elevated relative drag in high-frequency fliers arises primarily from increased drag coefficients.

Across species, weight-normalized aerodynamic power scaled positively with body mass as 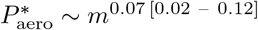 (PGLS regression: *R*^2^ = 0.18, *p* = 0.003; Table 1; Fig. 5A). Larger Diptera therefore require a higher relative power output during hovering flight. However, the observed PGLS slope (*a* = 0.07 [0.02 – 0.12]) was significantly lower than the theoretical scaling required for maintaining weight support across sizes (*a*_ws_ = 0.17) or as predicted under geometric, kinematic and dynamic similarity (*a*_sim_ = 0.33). This indicates that relative power scales negatively allometric, and that smaller Diptera experience higher relative power demands than predicted under similarity-based scaling.

Culicomorpha and Tipulomorpha are again striking outliers, with a relatively high and low aerodynamic power requirement, respectively. Importantly, these trends distinguish relative from absolute extremes. For example, Culicomorpha (mosquitoes and midges) exhibit elevated weight-normalized aerodynamic power relative to similarly sized Diptera, yet on an absolute scale they are not unusually high (e.g., compared with much larger droneflies).

According to our model (Eq. 6), the weight-normalized aerodynamic power scales with the product of wing speed and relative drag 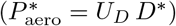.This relation allows us to separate kinematic effects (*U*_*D*_) from aerodynamic effects (*D*\*), and therefore identify the mechanisms underlying the elevated power requirements of tiny Diptera in general and Culicomorpha specifically. For this, we first validated the model by comparing the predicted power with CFD-based estimates. The two measures were strongly correlated (r = 0.92, p *<* 0.001), indicating that the simplified model (Eq. 6) captures the CFD-derived power expenditure. We then used a series of PGLS regressions to quantify the independent contributions of kinematics and aerodynamics to aerodynamic power demand and to identify the drivers of high power requirements in small Diptera and Culicomorpha (Fig. 5A,B,E).

The PGLS regression for wing speed shows that it scales positively with body mass as *U*_*D*_ ~ *m*^0.16 [0.13 – 0.19]^ (Fig 5B, Table 1). This scaling is significantly lower than expected under morphological and kinematic similarity (*a*_sim_ = 0.33), but closely aligns with the theoretical scaling required for weight support (*a*_ws_ = 0.17). Moreover, the mass-specific scaling of wing speed during hovering flight seems highly conserved among Diptera, including Culicomorpha with their high-frequency low-amplitude wingbeats (*U*_*D*_ ~ *f A*). The flapping wings of Tipulomorpha seem to operate at relatively low speeds, for their respective size.

The PGLS regression of relative drag shows that drag scaled negatively with body mass as *D*\* ~ *m*^−0.10 [−0.14 – −0.06]^ (Fig 5E, Table 1). This scaling deviates from the mass-independent scaling as assumed under dynamic similarity (*D*\* ~ *m*^0^), and is primarily caused by the elevated relative drag in tiny Diptera, and especially in Culicomorpha.

The relative drag is defined as the ratio between the drag and lift coefficients (*D*\* = *C*_*D*_*/C*_*L*_). The temporal dynamics of these CFD-derived coefficients reveal the mechanism underlying the elevated drag in Culicomorpha (Fig. 5C–D). Across most Diptera, the time-varying profiles of both *C*_*D*_ and *C*_*L*_ are highly conserved, but Culicomorpha form a clear exception. Their drag coefficients reach markedly higher values, particularly at the start of each wing stroke. As a consequence, the increase in relative drag in Culicomorpha arises primarily from elevated *C*_*D*_ rather than from reduced *C*_*L*_.

These results show that the weight-normalized aerodynamic power scales positively with body mass, but with a weaker size dependence than predicted under geometric, kinematic, and dynamic similarity (Fig. 5A). This reduced exponent arises because relative aerodynamic drag increases in the smallest Diptera (Fig. 5E). As a result, the expected decrease in wing speed with size is partly offset by an increase in drag at low Reynolds numbers. Tipulomorpha lie below the mass-scaling trend because they achieve markedly lower power requirements through reduced wing speeds (Fig. 5B). In contrast, Culicomorpha fall well above the trend because their high-frequency, low-amplitude wingbeats generate exceptionally high relative drag (Fig. 5E).

The rise in relative drag among small Diptera is consistent with expectations for insect wings operating in the intermediate Reynolds number regime [27, 28]. In this regime, the aerodynamic drag coefficient transitions between two asymptotic limits: an inertial high-Re regime where the drag coefficient is approximately independent of Reynolds number (*D*\* ∝ Re^0^), and a laminar–viscous low-Re regime where the drag coefficient increases as Reynolds number decreases 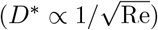. Consistent with this theoretical framework, relative aerodynamic drag scaled negatively with Reynolds number as *D*\* ~ Re^−0.20 [−0.28 – −0.11]^ (Fig. 6A; Table 1); this exponent lies between and is significantly different from both asymptotic predictions. As a consequence, relative drag is approximately constant for large Diptera operating near the inertial regime, but increases in the smallest species as they start to transition into the laminar–viscous regime where viscous forces become more dominant.

**Fig 6.**
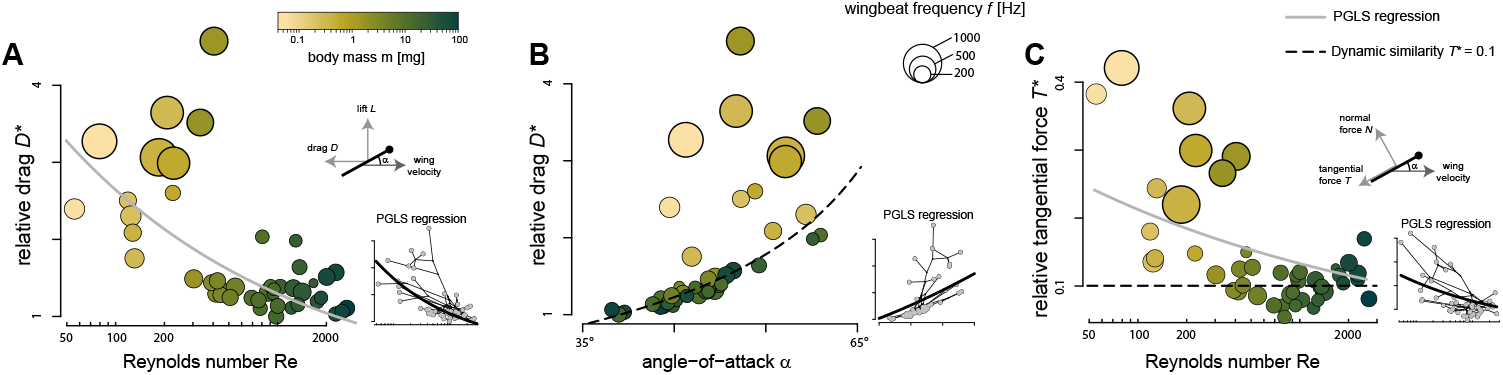
Scaling of drag with angle-of-attack and Reynolds number. In all panels, data points show species averaged results, color-coded by weight and size-defined by wingbeat frequency. Culicomorpha species with elevated wingbeat frequencies and swarm-based mating are highlighted in bold. Grey lines show PGLS regressions, and dashed lines show results for a constant relative tangential force (*T* * = 0.1). (**A**) The PGLS regression shows that relative drag (*D*\*) decreases with Reynolds number; high-frequency fliers exhibit the highest *D*\*. (**B**) For larger Diptera, *D*\* increases with angle-of-attack in accordance with force redirection at approximately constant relative tangential force (*T* * = 0.1; dot-dashed line). Small Diptera (*m <* 2 mg), and especially Culicomorpha, show much higher *D*\* than predicted by this trend. (**C**) The PGLS regression shows that the relative tangential forces (*T* *) decrease with Reynolds number; Culicomorpha experience exceptionally high *T* *.

Relative drag depends on both the relative (viscous) tangential force acting on the wing (*T* * = *C*_*T*_ */C*_*N*_), and the angle-of-attack of the wing (*α*). To control for this angle-of-attack effect, we correlated the CFD-derived relative drag with wingbeat-average angle-of-attack (Fig 6B). As expected, the relative drag increased with angle-of-attack (PGLS regression, Table 1), and most Diptera species closely align with the expected scaling for a constant relative tangential force *T* * = 0.1. In contrast, tiny Diptera (*m<*2 mg) lay above this constant-*T* * iso-line, suggesting that they experience increased tangential (viscous) drag. Hereby, the Culicomorpha are the greatest outliers.

We continued to test this by correlating the CFD-derived relative tangential forces with wingbeat-average Reynolds number at which the flapping wings operated (Fig 6C, Table 1). Again, the relative tangential force on the wings scale negatively with Reynolds number as *T* * ~ Re^−0.20 [−0.31 – −0.09]^, confirming that relative tangential forces on flapping Diptera wings scales inversely with Reynolds number. Like for relative drag, this inverse scaling lies in between and is significantly different from both asymptotic expectations for the intermediate Re regime (*T* * ∝ Re^0^ and 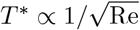). This confirms that the Re-dependent increase in relative drag on flapping Diptera wings is primarily caused by a viscosity-induced increase in relative tangential forces [27, 28]. It also shows that relative drag is rather constant for larger Diptera that operate near the inertial Re-regime, but rapidly increase in the tiniest Diptera due to a move into the laminar–viscous Re-regime.

These dynamics also explain why aerodynamic power scales more weakly with body size than predicted for maintaining weight support under dynamic similarity (Fig. 5). As Reynolds number decreases, tiny Diptera experience a disproportionately high viscosity-induced drag, which partly offsets the expected reduction in aerodynamic power associated with their lower wing velocities 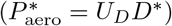. As a consequence, the balance between inertial and viscous contributions shifts progressively across the intermediate Reynolds-number range, moving the scaling of 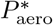 between the two asymptotic expectations. Culicomorpha clearly departs from the trend due to their unusually high drag.

### Scaling of muscle and aerodynamic power with size and phylogeny

To address the final component of our second objective (ii), the physiological requirement for flight muscles to deliver sufficient mechanical power, we compared the aerodynamic power required for flight with the relative muscle mass and expected power output of these muscles (Fig 7).

**Fig 7.**
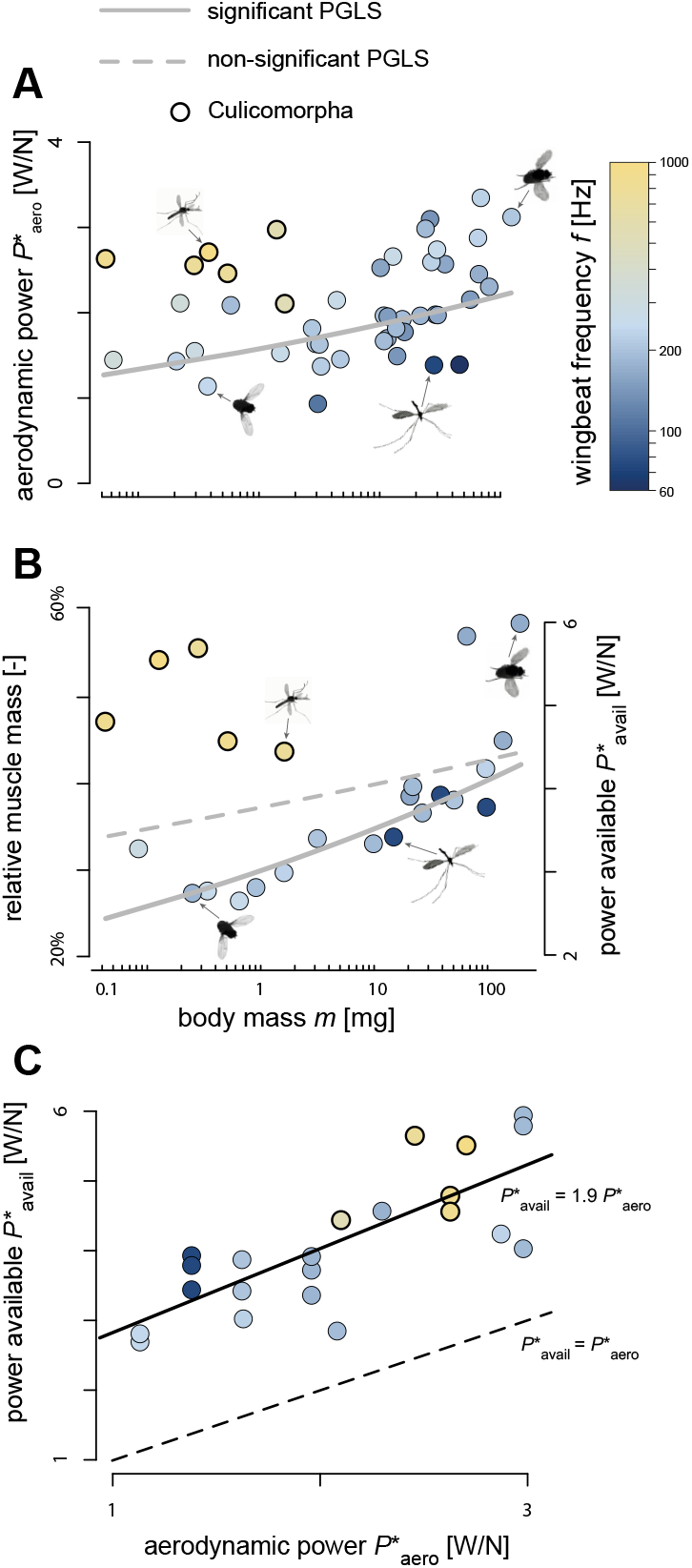
Scaling of required aerodynamic power and musculature-based available power. Data points show results per species, and are color-coded with wingbeat frequency (top right). Culicomorpha species with elevated wingbeat frequencies and swarm-based mating are highlighted in bold. (**A-B**) Aerodynamic power requirement (**A**, same panel as Fig 5A) aligns with the scaling of available flight power (**B**), estimated from thorax mass in a subset of 22 species. Data points in (**A**) represent species means, whereas data points in (**B**) represent individual measurements. Trend lines are based on statistical PGLS fits; the significant PGLS in (**B**) excludes Culicomorpha data. (**C**) Required aerodynamic power versus available power. The solid line shows the best OLS fit on the data, and the dashed line shows equal required and available power 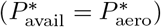. Independent of size, Diptera possess on average a two-fold higher available power than the aerodynamic power required for flight.

The available relative muscle mass did not show a significant scaling with body mass in all Diptera (PGLS regression: R^2^ = 0.12; p = 0.193), largely due to the Culicomorpha species possessing a markedly higher relative muscle mass than other taxa (Fig 1C and Fig 7B). However, when Culicomorpha were excluded, the relative muscle mass increased significantly with body mass, as 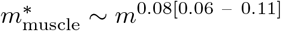 (PGLS regression: R^2^ = 0.85; p *<* 0.001). This scaling factor deviates significantly from the similarity expectation of ~ *m*^0^, and is strikingly similar to the the mass-specific scaling of the relative aerodynamic power for Diptera (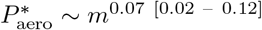; Fig 6A and Fig 7A). Moreover, the direct comparison between required aerodynamic power and the available muscle power shows strong similarities (Fig 7A and B, respectively). Among the larger Diptera both quantities increase with body mass; this dependence weakens toward smaller sizes and levels off in the tiniest species that operate near the laminar–viscous Reynolds-number regime.

Directly comparing the required and available power across species shows that Diptera consistently possess approximately twice the power required for flight (Fig 7C). This relationship is consistent across taxa, including both the energetically inefficient Culicomorpha and the more efficient Tipulidae.

### Scaling of acoustic power with size and phylogeny

To investigate if lineage-specific selective pressures promote departures from energy-minimizing flight (Objective iii), we analyzed the scaling of acoustic power across the phylogeny. Specifically, we tested whether and how the high-power wingbeat of mosquitoes and midges produce increased aero-acoustic sound, potentially driven by sexual communication. For this, we estimated the far-field sound produced by the beating wings by augmenting our CFD results with aero-acoustic modeling (Fig 8). This shows that sound power scales positively with body mass as *P*_acoustic_ ~ *m*^1.57[1.38 – 1.76]^ (PGLS regression: R^2^=0.86; p *<* 0.001; Fig 8A). This scaling is not significantly different from the expected scaling for maintaining weight support under geometric similarity (*P*_acoustic_ ~ *m*^5*/*3^; see Supplementary Methods S1.8 for details). For Diptera in general, there thus seem no allometric adaptations present for sound production. In contrast, this is clearly different for the infraorder Culicomorpha, who exhibited disproportionately high sound power levels for their small size (Fig 8A). Within this size-controlled, phylogenetically informed analysis, species that incur elevated aerodynamic costs for their size also tend to exhibit elevated aero-acoustic power. Thus, prior qualitative statements about a flight–acoustics linkage are here converted into tested comparative trends, with the strongest divergence observed in Culicomorpha.

**Fig 8.**
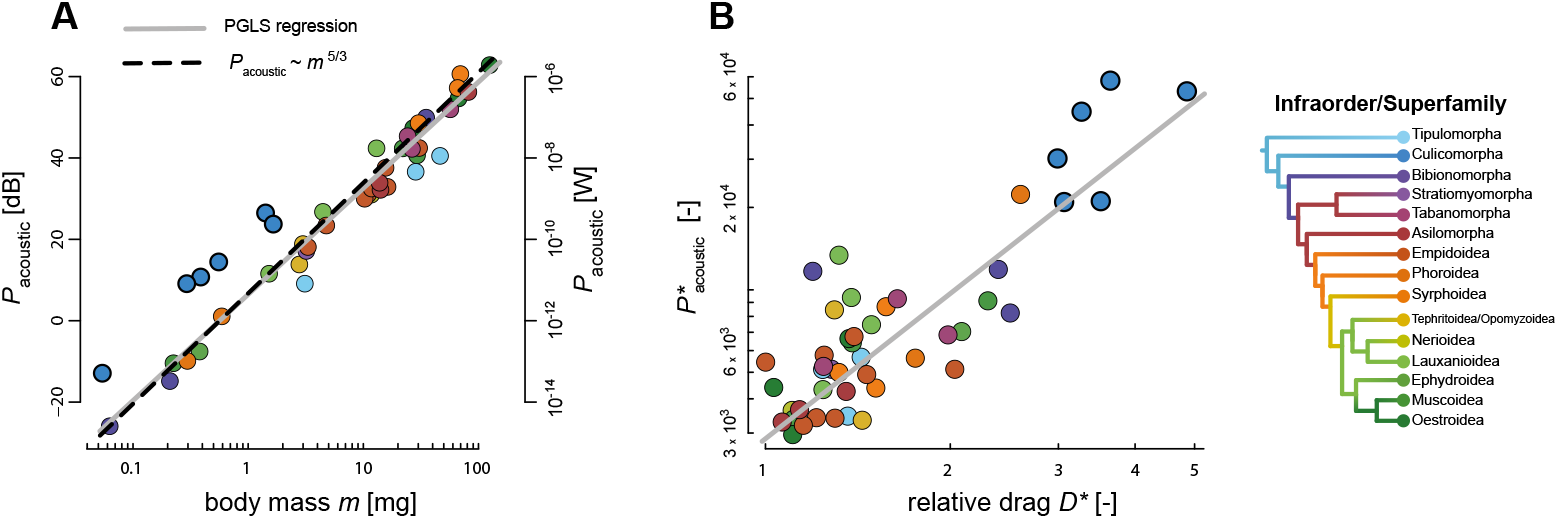
Scaling of aero-acoustic power with body mass and aerodynamic drag production. Data points show species-average results, and are color-coded by infraorder (see right). Culicomorpha species with elevated wingbeat frequencies and swarm-based mating are highlighted in bold. The gray and black lines show significant PGLS regressions and theoretical scalings, respectively. (**A**) Aero-acoustic power increases with body mass across Diptera, with Culicomorpha species exhibiting enhanced acoustic power compared to other Diptera of similar sizes. The PGLS regression closely matches the theoretical scaling expectation *P*_acoustic_ ~ *m*^5*/*3^. (**B**) Normalized acoustic power 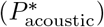 scales positively with the relative aerodynamic drag (*D*\*), whereby Culicomorpha produce both highest normalized sound power and relative drag.

Secondly, we estimated the normalized aero-acoustic power 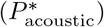 from CFD, kinematics, and morphology (Eq. 9), and tested how it scales with relative drag *D*\* (Fig. 8B). The resulting PGLS regression was positive, showing that acoustic power and aerodynamic power increase simultaneously, consistent with a trade-off between sound production and aerodynamic cost minimization. Notably, Culicomorpha did not break this trend; instead, these species exhibited both elevated acoustic power and elevated relative drag compared with other Diptera. This indicates that Culicomorpha generate their enhanced acoustic output primarily by producing increased relative aerodynamic drag.

## Discussion

Here, we present one of the most comprehensive comparative biomechanical studies of insect flight within a phylogenetic framework, investigating the factors influencing flight diversity in Diptera, one of the largest and most ecologically diverse insect orders [1, 41, 42]. We quantified wingbeat kinematics, morphological traits, and aerodynamic force and power production, and aero-acoustic sound production across the phylogeny, thereby revealing how biomechanical traits diversifies across Diptera. By evaluating the extent to which the observed diversity in these traits can be explained by physical scaling laws and aerodynamic principles, we also highlight deviations that likely reflect adaptive responses to selective pressures.

A central contribution is that we place well-known single-taxon observations (e.g., slow flapping in crane flies, enlarged thoracic musculature in robber flies, high-frequency low-amplitude wingbeats in mosquitoes) into a unified, phylogenetically controlled, size-aware framework. This allows us to quantify how extreme a lineage is relative to similarly sized Diptera, to separate relative from absolute extremes (e.g., aerodynamic cost in mosquitoes), and to test linkages between aerodynamic costs and acoustic power as comparative patterns rather than species-specific observations. Moreover, our comparative framework is anchored in full-DNS CFD simulations, which provide high-fidelity aerodynamic force and power estimates and inherently resolve induced power, wake interactions, and other unsteady aerodynamic mechanisms unique to insect flight; this ensures that all comparative inferences account for the complete aerodynamic cost structure of flapping flight.

To support further comparative and evolutionary analyses, all morphological, kinematic, and aerodynamic datasets used in this study is available in an online repository. Here, we also provide a tool in which the dataset can be interactively explored and downloaded, enabling users to examine clade-specific patterns and trait relationships beyond the static figures presented here. See the Data Availability Statement for details.

### Conserved and divergent traits in Diptera flight motors systems

Our comparative analysis of the Dipteran flight motor system sheds light on the relative contributions of kinematic and morphological diversity to the evolution of Diptera flight (Objective (i)). Despite the wide ecological diversity of Diptera and its potential influence on trait evolution, our results show that wingbeat kinematics is relatively well conserved across their phylogeny. Even among highly divergent lineages, flight is achieved through remarkably similar wingbeat kinematics, with conserved temporal dynamic of wing stroke, deviation, and rotation angles. This demonstrates the dominant role of aerodynamic constraints in shaping the evolution of flight. The persistence of such a broadly consistent flapping-wing propulsion system across diverse Dipteran taxa, and potentially more generally across insects and even other flapping-flying animals [4], suggests that it may have been key to their ability to exploit a wide range of ecological niches [42]. Moreover, this pattern shows that similar wingbeat kinematics can arise from a wide range of wing and body morphologies, suggesting a many-to-one mapping of form to function as described in other biomechanical systems (e.g., [73, 74]).

Two notable exceptions stood out from this general pattern: species from the infraorders Culicomorpha (mosquitoes and midges) and Tipulomorpha (crane flies). In Culicomorpha, wingbeat frequency was exceptionally high and stroke amplitude markedly reduced compared to other Diptera. Conversely, Tipulomorpha species exhibited markedly low wingbeat frequencies and angular wing speeds, relative to all other taxa. This suggests that flight in these two early-diverged lineages may have been shaped by distinct selective regimes compared to more recently diverged taxa. For clarity, references to early diverged taxa pertain to extant lineages and should not be interpreted as statements about the phenotype of ancestral Diptera.

Variation in wingbeat kinematic was nonetheless affected by phylogeny. Indeed, we detected a significant phylogenetic signal in most wingbeat kinematic parameters, except for wing angle-of-attack, which showed very limited interspecific variation and thus appears to be the most tightly constrained by aerodynamic requirements.

In comparison with wingbeat kinematics, variation in wing and body morphology across species was more structured by phylogeny. This finding challenges the general view that behavioral traits are more evolutionarily labile than morphological traits [60]. If kinematics were as labile as generally assumed for behavioral traits, the diversification of Diptera species may have resulted in more diverse wingbeat kinematics, possibly driven by adaptations to specific ecological niches, even among morphologically similar species. The fact that kinematics appear so conserved across species further emphasizes the tight constrained exerted by aerodynamic requirements, which in turn may have favored adaptations to occur through morphological rather than kinematic changes among Diptera.

Our results show that morphological shape (including wing and body aspect ratio) primarily distinguishes Dipteran taxa, with the earliest diverged lineages exhibiting elongated high aspect-ratio forms, and recently diverged ones showing stouter low aspect-ratio morphologies. Whether evolutionary changes in wing and body shape across Diptera are primarily adaptive or stem from neutral evolutionary processes is challenging to ascertain. Early-diverged Diptera might have retained morphologies closer to the ancestral state (putatively slender body and high aspect-ratio wings), while the diversification of subsequent lineages into varied ecological niches may have promoted the evolution of stouter wing and body shape. Such morphology reduces the moment of inertia, allowing faster rotations and overall enhancing maneuverability – a key component of flight performance under strong selection for tasks such as capturing prey, evading predators, or navigating cluttered environments [13, 75, 76]. This advantage, however, may come at the cost of increased energetic demands. A possible interpretation to the observed morphological pattern is that selective pressures favoring enhanced maneuverability were less pronounced during the diversification of early-diverging lineages compared to the more recently diverged groups [77].

Assessing the adaptive significance of these trait changes is particularly difficult at the order-wide scale of our study. Indeed, adaptive divergence is more readily inferred when comparing closely related species with contrasting ecologies (e.g., [30, 78, 79]), where phylogenetic inertia and unknown selective pressures play a smaller role. While our macroevolutionary analysis reveals broad patterns, finer-scale comparative work is needed to disentangle neutral divergence from ecological adaptation. In contrast with morphological shape, body size was more variable within taxa, suggesting that size might be less phylogenetically constrained than shape in Diptera. Body size, however, imposes strong physical and aerodynamic constraints on flight-related traits.

### Scaling and aerodynamic constraints structure Diptera flight diversity

Much of the variation in morphological and biomechanical traits can generally be explained by scaling laws, i.e., how traits change with body size [80–82]. In flying animals, an additional key constraint is the need to generate sufficient lift to support body weight, which strongly affect the evolution of morphological and biomechanical traits [3, 6]. To address Objective (ii), we tested how these constraints shape the scaling of morphology, kinematics, and aerodynamics across the Diptera phylogeny.

#### Weight support requirement in hovering flight

Our analysis of size–trait scaling shows that physical scaling laws and aerodynamic constraints strongly structure Diptera flight. For size reductions under geometric similarity, wings generate proportionally less aerodynamic force, which requires compensatory changes in morphology or kinematics. Consistent with this expectation, we found that smaller species exhibit relatively larger wingspans and slightly higher normalized radii-of-gyration, placing more wing area distally and resulting in increased flapping-wing-based force generation. Wingbeat kinematics also change with size: wingbeat-induced angular speeds increases markedly in smaller species, primarily caused by an increase in wingbeat frequency; wingstroke amplitude remains approximately size-independent. Also angle-of-attack shows a weak negative allometry with size, but this reduces the wingbeat-average force coefficient for tiny Diptera.

Together, these allometric adjustments increase aerodynamic force production capacity as size decreases, and therefore compensate for the reduced lift expected under geometric similarity. Comparing the fitted scaling exponents to similarity predictions and to those required for weight support showed that wingbeat-kinematic adjustments provide the dominant contribution, whereas allometric increases in wing size form a secondary contribution (*a*\* = 84% and 51%, respectively). In contrast, adaptations in wing shape, expressed by the normalized radius of gyration, contribute only modestly (4%). The angle-of-attack effect opposes compensation because it lowers the wingbeat-averaged force coefficient, resulting in a negative contribution (*a*\* = –11%).

Note that the sum of all significant relative allometric scaling factors exceeds 100% (128%). This suggests that different taxa exhibit distinct kinematic and morphological adjustments, and that our analysis captures their combined effects, thereby overestimating the order-wide pattern. This is most apparent from the influence of outlier groups: the extremely high wingbeat frequencies in small Culicomorpha might inflate the frequency scaling; in contrast, Tipulidae show wingspan scaling close to geometric similarity, which might reduce the apparent size dependence of wingspan across Diptera.

Taken together, our scaling analysis shows that Diptera use both kinematic and morphological adjustments to maintain weight support across body size, albeit potentially using distinct mechanisms among different taxa. These results are in line with patterns observed in other flying animals, where in some cases kinematic adaptations dominate [5, 6, 38, 83–85], and in others morphological compensations prevail [20, 86–89].

#### Power requirement for hovering flight in Diptera

Our Computational Fluid Dynamics (CFD) simulations show that aerodynamic power scaling in Diptera follows expectations for wings operating in the intermediate Reynolds number (Re) regime. In this regime, aerodynamic drag transitions between two asymptotic limits: an inertial high-Re limit where the drag coefficient is approximately Re-independent, and a laminar–viscous low-Re limit where the drag coefficient increases as Re decreases [27, 28, 90]. Classical Rankine–Froude hovering theory aligns with the high-Re inertial limit and predicts that mass-specific induced power scales as 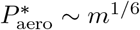 under geometric similarity [26].

In our dataset, the CFD-derived weight-normalized aerodynamic power approaches this inertial expectation in larger Diptera, where relative drag is effectively size-independent and wing speed scales near *U*_*D*_ ~ *m*^1*/*6^. Toward small sizes (*m <* 2 mg), the size dependence of 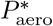 weakens. This shift reflects the Re-dependent rise in relative aerodynamic cost: as Re declines, viscous effects amplify relative tangential forces (*T* *) and consequently increase relative drag (*D*\*) [28, 90, 91]. This increase persists after accounting for variation in mid-stroke angle-of-attack, indicating a genuine viscosity-driven effect rather than a kinematic artifact. Because 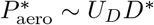, two opposing trends counteract each other with decreasing size: the decrease in *U*_*D*_ lowers aerodynamic power requirement for flight, while the increase in *D*\* raises it. The net result is a muted size dependence of 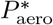 among the smallest Diptera, consistent with an approach toward the laminar–viscous asymptote.

Thus, for insects with body masses below ~2 mg, further miniaturization faces a progressive physical constraint. The viscosity-induced increase in drag limits additional gains in aerodynamic economy, even though absolute aerodynamic power remains small. This pattern does not imply a hard lower size limit for powered flight in Diptera, but reflects that small Diptera pay an increasingly viscous cost for lift production along the inertial-laminar–viscous continuum. Beyond Diptera, viscosity may limit flight performance in the smallest specialized fliers that operate truly in the laminar–viscous Reynolds-number regime [28, 29, 91, 92].

At the opposite end of the Diptera size range, hovering becomes more demanding because absolute aerodynamic power rises steeply with body mass, making sustained hover a stronger constraint for large Diptera [3, 10, 26].

The scaling of available power for flight, inferred from thorax mass, closely matched the scaling of aerodynamic power requirements from CFD simulations, underscoring the tight co-evolution of muscle morphology and aerodynamic demands [26, 91]. This alignment ensures that power supply and demand remain balanced, avoiding performance failure or metabolic inefficiency [3, 10]. We show that hereby most Diptera maintain a two-fold safety margin in power availability, consistent with previous findings [56, 70]. This excess muscle capacity likely safeguards performance under demands for maneuverability or environmental variability (e.g., predator attack or windy conditions, respectively). This consistent safety margin even includes Culicomorpha and Tipulomorpha species, in which we observed an increase and reduction of both power requirements and flight musculature, respectively. The increased musculature in Culicomorpha enables them to fly with their exceptionally costly high-frequency wingbeats. These results highlight how physical constraints, energetic optimization, and lineage-specific selective pressures jointly shape the evolution of the Diptera flight motor system.

The observed scaling between aerodynamic power requirements, available motor power, and body mass provides a baseline around which most Diptera species cluster. Deviations from this trend may point at evolutionary trade-offs. While insufficient aerodynamic power risks flight failure, excessive investment in increased flight power musculature may reflect selective pressures favoring specialized flight performance. For example, higher power demands may be under selection in species where rapid flight acceleration is crucial for survival or reproduction, such as predatory insects (e.g., robber flies [13]), or in species that engage in sustained aerial mate pursuits (e.g., hoverflies [19, 20]). By contrast, in species with lifestyles that place little demand on maneuverability, selection may favor reduced flight costs, as observed here in Tipulomorpha, where exceptionally low energetic demands may have evolved at the expense of maneuverability.

Strikingly, our results reveal a clear deviation from the general scaling trend in a subset of the smallest species. Culicomorpha (mosquitoes and midges) show power requirements nearly three times higher than other taxa of comparable size. Their elevated flight cost arises because their exceptionally high wingbeat frequencies generate highly unsteady aerodynamic forces [35, 93]. These increased flight costs are associated with enlarged relative thoracic muscles, necessary to sustain the required power output.

### Potential trade-off between aerodynamic and acoustic efficiency in mosquitoes and midges

The flight of mosquitoes and midges has been extensively studied in the context of sound production and its role in locating sexual partners in so-called mating swarms [21–23, 65, 94, 95]. Our third Objective (iii) was to investigate whether such lineage-specific selective pressures, like acoustic signaling, can drive a departure from energetic cost minimization. It has been shown that male and female mosquitoes adjust their flight tones during swarming, enabling acoustic mate recognition [24]. Selection for effective acoustic signaling, essential for mosquito reproduction, may thus come at the expense of flight cost minimization.

Consistent with the hypothesis that sexual communication imposes strong selective pressures on mosquito flight, our comparative analysis highlight the strong divergence of muscular morphology and wingbeat kinematics between Culicomorpha and all other studied Diptera. Moreover, we showed that mosquitoes and midges produce substantially higher acoustic power for their size compared to other Diptera, which they achieve by simultaneously increasing their wingbeat frequency and reducing their wingstroke amplitude. These adaptations in wingbeat kinematics come at the expense of an increase in aerodynamic power requirement for hovering flight, and a corresponding increase in flight motor musculature.

These combined results suggest that Culicomorpha have followed an evolutionary trajectory shaped by strong selection on acoustic signaling during swarm-based mate localization. Their extremely high wingbeat frequency, low stroke amplitude, and elevated aerodynamic and acoustic power are consistent with selection for effective flight-tone production and detection. As a consequence, these traits point to an evolutionary compromise in which reproductive signaling demands impose increased aerodynamic cost, contributing to the marked divergence of Culicomorpha from the kinematic and aerodynamic patterns observed in other Diptera.

### Scope and limitations of the study

Our Culicomorpha dataset consists of male-only singular single-individual flight recordings. In many Culicomorpha, acoustic interactions during in-flight courtship are actively coupled: males detect female flight tones and adjust their wingbeat to maintain interference (difference) tones (e.g. [21–24]). Because our dataset includes male individuals only, and because single-individual flight recordings cannot reproduce the interactive acoustic loop that occurs during paired flight, we interpret the elevated aerodynamic and aero-acoustic output in Culicomorpha as a plausible consequence of sexual selection rather than a definitive explanation. Additional ecological pressures, such as load carrying after blood meals [55], may also contribute to the observed divergence. Demonstrating causal mechanisms will require dedicated future experiments with sex-balanced sampling, paired co-flight measurements, and auditory modeling.

Next to Culicomorpha, male and female individuals may also exhibit divergent flight behaviors in other Diptera taxa [18, 19, 96], although such differences are expected to be more subtle and therefore remain poorly documented. Because we did not record the sex of individuals among the non-Culicomorpha species, our comparative analysis cannot address sex-specific differences in flight. We therefore did not include this dimension of flight diversity into our analyses, and assume that interspecific variation across the order exceeds intraspecific sexual differences in flight behaviour.

Our aerodynamic force and power estimates represent the *total* aerodynamic force and power produced during hovering flight, because they are computed directly from full-DNS CFD. These totals include induced, viscous, and unsteady contributions (including wake-mediated mechanisms such as wake capture) in an integrated manner. However, the present comparative dataset is not designed to provide a mechanistic decomposition of these totals into components such as lift-induced power, viscous drag-induced power, or other unsteady aerodynamic terms.

Likewise, our analysis focuses on aerodynamic forces only, and therefore inertial forces associated with accelerating and decelerating the wings are not included. Including these additional contributors to overall flight force and energetics would require further measurements and modeling, which lie beyond the scope of this order-wide comparative study. We therefore treat these topics as priority directions for future work enabled by the openly available morphological, time-resolved kinematic, and aerodynamic datasets.

Next, all aerodynamic forces and power estimates in this study are derived from near-hovering flight, which we use as a standardized comparative benchmark. This choice facilitates cross-species comparison but does not capture performance during forward or maneuvering flight.

Finally, our estimates of available flight power are based solely on flight-muscle mass. Mechanical power output depends on additional traits that we did not quantify, including muscle fibre architecture and the mechanical coupling between muscle and thorax [70, 71, 97, 98]. These anatomical and physiological properties vary across taxa and may influence both the magnitude and the scaling of available power. Future comparative work should therefore target these components of the flight motor system to better understand how muscular specialization contributes to divergence in flight performance across Diptera.

### Broader comparative context and conclusions

The patterns identified here have implications extending beyond Diptera. Size-dependent aerodynamic limits on hovering, where absolute power requirements rise steeply with body mass, are expected in other insect groups, and in any flapping flier operating in the intermediate Reynolds number regime. Similar constraints have been documented across insects and vertebrates, including classical analyses of induced power in flapping birds and insects [3, 10, 26]. The conserved wingbeat kinematics observed in Diptera parallels these broader biomechanical patterns and suggests that flapping-flight systems across animals face similar physical boundary conditions.

Conversely, taxa-specific deviations, as seen in Culicomorpha, illustrate how ecological or sexual selection can drive departures from these constraints. Comparable divergences have been described in other flying animals, for example in birds [99, 100], moths [79], or dragonflies [101], where specialized ecology or signaling demands shift morphology and kinematics away from energy cost minimization. Such deviations generate testable hypotheses about how selective pressures may repeatedly reshape the flight motor system across animal lineages.

Overall, our comparative analysis shows how physical scaling laws, aerodynamic constraints, and ecological or sexual selection combine to structure the evolution of complex locomotor systems. The integrative framework developed here provides a mechanistic basis for interpreting flight diversity within Diptera and offers a general template for future comparative studies across flying animals, with relevance for evolutionary biology, functional morphology, and bio-inspired design.

## Acknowledgments

The authors gratefully acknowledge the late Paul Beuk, whose enthusiasm and knowledge of Diptera taxonomy left a lasting impression and provided valuable context for this work. We thank Antoine Cribellier, Guillermo Amador, and Abel-John Buchner for helpful discussions on the quantification of insect flight. We are grateful to Remco Pieters for his assistance in building and operating the experimental setup, and to Henk Schipper and Michel Breuer for their support with laboratory work. We thank Serge Poda and Tessa Visser for providing the lab-reared Diptera specimens, and Uroš Cerkvenik for his help in determining relative muscle data of large horseflies.

## Data Availability Statement

All data underlying the findings of this study are fully available without restriction. The complete datasets used in the analyses—including wing morphology, wingbeat kinematics, and CFD-derived aerodynamic forces and power, as well as CFD-parameter files, microscope images and representative high-speed videos—are deposited in the Dryad Digital Repository under DOI: 10.5061/dryad.gxd25480s. Here, we also provide an interactive tool with which users can filter, visualize, and export all processed data, or subsets of these. DNA barcode data for all analyzed specimens are publicly available through the BOLD System under project DIPTR via this link. No ethical or legal restrictions apply to the sharing of these data.

## Supporting information

**S1 Text - Extended Methods Extended materials and methods**. This document provides the extended experimental and analytical framework used in our study, including specimen sampling, morphological and kinematic measurements, flight-experiment procedures, and CFD simulations. It also contains the aerodynamic model based on Buckingham–Pi dimensional analysis, the scaling relations for force, power, and aero-acoustic output, and additional details on how morphological and kinematic traits jointly contribute to maintaining weight support across sizes.

**S1 Fig. Estimation of fresh body mass**. The fresh body mass of specimens collected in French-Guiana was approximated using the relationship between dry and the fresh body mass among specimens collected in Wageningen. (**A**) A linear regression model was fitted to the data from Wageningen. (**B**) The equation of the fitted model was used to estimate fresh body mass: *m* = 4.1 ∗ *m*_dry_ − 0.96.

**S2 Fig. Intra-specific and inter-specific flight variation**. Intra-specific versus inter-specific variation in flight kinematics parameters assessed with the coefficient of variation (*C*_var_). Parameters are ordered by decreasing inter-specific *C*_var_.

**S3 Fig. Comparison between normalized radius of gyration 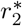 and center of the aerodynamic drag force 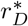.**

**S1 Video. High-speed flight recordings of the 46 Diptera species analyzed**. Collage of high-speed videos of all 46 Diptera species studied in flight. The separate videos have been scaled and aligned such that they fit the collage and video duration, and thus spatial and temporal scales differ between them. The corresponding full videos are available in the digital repository (see Data Availability Statement). All videos were recorded and edited by the authors.

**S1 Table. Phylogenetic signal in morphological and wingbeat kinematic parameters, computed using Blomberg’s K**. Bold indicate significant p-values (p*<*0.05).

**S2 Table. Comparison of sum of squares from nested ANOVAs accounting for multiple flight sequence replicates per species**.

