## Supplementary material for "Diptera flight diversity is shaped by aerodynamic constraints, scaling, and evolutionary trade-offs": S1 Text

### S1 Text – Extended Methods

#### *S1.1. Insect sampling*

We studied the diversity of Diptera flight motor systems by sampling 133 species distributed across 43 families that together span the phylogeny and size range of the order (Fig. 1A,B). Most material (94 species) was collected in the surroundings of Wageningen University, the Netherlands (51°59′01.0″N, 5°39′32.7″E; ~10 m a.s.l.) during the summers of 2021 and 2022. Additional sampling in the Amazonian rainforest of French Guiana (4°34′09″N, 52°13′04″W; ~300 m a.s.l.) in July 2021 yielded 36 species. Three mosquito species were obtained from laboratory cultures. These collections yielded a mean ± standard deviation of 3.2 ± 3.7 species per family and 1.9 ± 1.2 individuals per species.

Wild-caught individuals were identified using morphology and DNA barcoding. For barcoding, we submitted a leg from each individual to the Canadian Centre for DNA Barcoding (CCDB), where barcodes were generated using the standard automated protocols of the BOLD Identification System. Resulting sequences were matched against the BOLD reference libraries to establish species-level identifications when possible. The phylogenetic context followed the comprehensive Diptera molecular phylogeny of Wiegmann et al. (2017)[1], which we pruned to the set of species included in our analyses (Fig. 1B).

Sample sizes per species were generally insufficient for sex-specific analyses, and we therefore did not determine sex for most taxa. An exception was made for Culicomorpha (mosquitoes and midges), for which we systematically analysed males only because males form swarms and are the phonotactically active sex that detect female flight tones and the interference (difference) tone generated during close-range interactions [2–4]. This male-specific behavior ensures biological relevance for aero-acoustic comparisons within this lineage but limits sex-specific inference.

Downstream datasets derived from this sampling include external body and wing morphology for all species, flight-muscle mass estimates for a subset of 23 species chosen to span phylogeny and size, and hovering flight kinematics with corresponding computational fluid dynamics (CFD) for 46 species chosen for broad taxonomic and size coverage (Fig. 1A,C). For figure clarity we group species by major lineages (infraorders or superfamilies), while all phylogenetic comparative analyses are performed at the species level.

#### *S1.2. Body and wing morphology estimation and shape analysis*

To quantify body and wing morphology, individuals were first euthanized by freezing them at −20 °C. Masses of the animals and their body parts were measured using analytical balances. For insects and body parts larger than a fruit fly ($m > 0.3$ mg), we used a balance with a resolution of 0.1 mg (XSR204 Mettler Toledo), while for smaller insects the balance had a resolution of 1 µg (Sartorius M2P). All measurements were performed within an hour after capture. Fresh body mass could not be directly measured for flies collected in French-Guiana. Therefore, we determined the relationship between dry and fresh body mass among all specimens collected in Wageningen as $m=4.1\cdot m_{dry}-0.96$ (S1 Fig). We then estimated the fresh body mass of the specimens from French-Guiana based on dry body masses using this relationship.

We quantified flight muscle mass on a subset of 23 species selected to span a broad phylogenetic and size range (Fig. 1A,C). For this, we dissected 5.0 ± 1.2 thoraxes per species and weighed them immediately after separation from the body. The thorax of flying insects mainly contains flight muscle, which determines the power output for flight [5]. Muscle mass was thus estimated from the thorax mass as $m_{muscle}=0.9\cdot m_{thorax}$ [6].

Individual flies and their detached wings were photographed under a digital microscope (Zeiss Stemi SV11). From these photographs, body length and thorax width were first measured using ImageJ (Schindelin 2012), and the body aspect ratio was computed as the ratio of body length to thorax width. Body length was measured along the mid-sagittal axis, from the anterior tip of the head to the posterior tip of the abdomen. Thorax width was measured as the maximum lateral width of the mesothorax in the horizontal plane, perpendicular to the body's longitudinal axis. We processed the wing images using *WingImageProcessor* in MATLAB (available at http://www.unc.edu/~thedrick/), which provided estimates of the following wing morphology parameters (Fig. 2A): wing surface area $S$, wingspan $b$, and the wing second moment of area relative to the wing hinge $S_2$. From these, we calculated mean wing chord as $\bar{c}=S/b$, wing aspect ratio $AR=b/\bar{c}=b^2/S$, the wing radius of gyration as $r_2=\sqrt{(S_2/S)}$, and the span-normalized radius of gyration as $r_2^*=r_2/b$.

We quantified differences in wing shape among species in more detail using a landmark-based geometric morphometric approach [7]. The outline of each wing was described with 300 semi-landmarks, equidistantly spaced along the wing outline. Semi-landmarks are used when identifiable landmarks are unavailable, typically to quantify curved shapes. To align the wings, the semi-landmarks were slid along the curve to establish geometric correspondence by minimizing bending energy [8]. After this alignment, the semi-landmarks were treated as regular ones in subsequent analyses. We did not consider vein architecture in the analysis because the large phylogenetic scale of our study limits the number of homologous identifiable vein points between species. Nonetheless, one fixed landmark was placed at the wing hinge to fix the overall landmark configuration relative to this homologous position available for all species. All landmarks were digitized using TpsDig2 [9]. Wing outlines were superimposed by performing a generalized Procrustes analysis [10], using the *geomorph* R package [11]. Through this process, shape information is isolated by discarding the variation in size, position, and orientation. To analyse and visualize wing shape variation between the sampled Diptera species, we then performed a principal component analysis (PCA) on the superimposed coordinates. The evolutionary structure of Diptera wing shape diversity was visualized by superimposing the phylogenetic tree onto the morphospace (Fig. 2A).

#### *S1.3. Flight experiments*

We studied the flight of 46 Diptera species covering 30 families, all collected in the surroundings of Wageningen University. Following capture in the wild, the insects were brought into the laboratory for the flight experiments. The experiments were performed in a custom-built octagonal flight arena with transparent Plexiglas walls (50×50×48 cm; height × width × length), adapted from Cribellier et al (2022) [12]. Three synchronized high-speed cameras (Photron FASTCAM SA-X2; spatial resolution 1024×1024 pixels) were positioned around the arena to provide a top view and two inclined side views. All cameras were aimed at the centre of the arena, where their focal planes intersected. Illumination was provided by infrared backlighting using high-power infrared LED panels (Osram OSLON Black Series 850 nm, 150°) placed behind the bottom and lower side walls opposite each camera.

We used two videography configurations, a zoomed-in and a zoomed-out configuration for smaller and larger insects, respectively. For species with body lengths larger than approximately 3 mm, the cameras were equipped with Nikon Sigma 135 mm lenses, resulting in a focal zone of approximately 12×12×12 cm in which the insect was in focus in all three views. For smaller species, the cameras were equipped with Canon EF 100–400 mm lenses paired with a 2× extender (effective focal length 200–800 mm), resulting in a focal zone of approximately 6×6×6 cm.

Camera frame rate was adjusted to the wingbeat frequency of each species. We aimed to record at least 20 frames per wingbeat to allow reconstruction of the full wingbeat kinematics. We used a minimum frame rate of 5000 frames per second for low-frequency flyers such as crane flies, and the maximum achievable frame rate of 13,500 frames per second for high-frequency flyers such as mosquitoes.

Before each experiment, the stereoscopic camera system was calibrated using the Direct Linear Transformation (DLT) [13]. Calibration was performed using a T-shaped wand with one ball at each end of the crossbar that was manually moved within the focal zone. The distance between the two balls was 2.8 cm in the zoomed-out configuration and 1.0 cm in the zoomed-in configuration. Ball positions were tracked using a neural network trained with *DeepLabCut* [14], and the DLT coefficients were calculated using the MATLAB program *easyWand* [15].

During each experiment, several individuals of the same species were released simultaneously in the arena to increase the probability of an insect flying through the focal zone. When this occurred, the camera system was manually triggered using an end-trigger to store the preceding video buffer. Filming continued until at least three flight sequences were captured per species. Because individuals could not be identified, it was not possible to assign flights to specific individuals. We therefore focused on determining average wingbeat kinematics per species. After the flight experiments, all insects in the arena were euthanized and their morphological characteristics were measured following the method described in the previous section.

#### *S1.4. Body and wingbeat kinematics reconstruction*

We quantified the body and wingbeat kinematics of all recorded flight sequences using an adaptation of the manual stereoscopic video tracker *Kine* in MATLAB, originally developed for tracking the wingbeat kinematics of flying fruit flies [16,17]. Here, we adapted the tracker by building wing models matching the wings of each studied species. These wing models were constructed by digitizing the wing outline from digital microscope photographs, resulting in 41 two-dimensional coordinates representing the shape of the wing outline, including the known hinge and tip positions. The tracking method is based on a rigid flat wing model, and thus wing deformations were ignored.

The body and wingbeat kinematics were tracked on videos comprising 34.6 ± 7.7 frames per wingbeat. To speed up digitization, videos were down-sampled for low-frequency fliers, to 2500 frames per second at its lowest. Through tracking, we acquired the position and orientation of the body and wings in each consecutive video frame, using the conventions described in Muijres (2023) [18](Fig. 3A). The body position was defined in the world reference frame as the temporal dynamics of the three-dimensional position vector **X**=[*x, y, z*], where the *z*-axis points vertically upward; body orientation was defined using the consecutive Euler angles body yaw ψ, body pitch β, and body roll ϕ, in the world reference frame. The angular orientations of the wings throughout the wingbeats were expressed in the stroke-plane coordinate system using three consecutive Euler angles (Fig. 3A) [18]: stroke angle ϕ, defined as wing-hinge rotations within the stroke-plane; deviation angle η, defined as rotations out of the stroke-plane; and the rotation angle θ, defined as wing rotations along the spanwise axis (Fig. 3B).

The stroke-plane was defined as a plane fixed within the body reference frame, located at the wing-hinge positions, and within which the majority of wingbeat motions were executed. Because the wing deviation angle is defined as angular rotation out of this plane, the stroke-plane is defined as the plane for which the deviation movements are minimal. The stroke-plane orientation is characterized by its pitch angle relative to the horizontal. We measured this stroke-plane pitch angle $\beta_{stroke\text{-}plane}$ by estimating the pitch angle for which the wingbeat-average deviation angle was minimal, using a Nelder–Mead optimization algorithm (Gao 2012) from the SciPy Python package.

The wingbeat kinematics were tracked for the left and right wings separately, and then averaged between the two wings. The time history of the averaged wing kinematic angles was then fitted with a fourth-order Fourier series for all three angles [19]. From this, we computed the wingbeat frequency as $f = 1/\Delta T$, where $\Delta T$ equals the time between consecutive wingbeats, and wingstroke amplitude as $A_\phi=\phi_{max}-\phi_{min}$, where $\phi_{max}$ and $\phi_{min}$ are the maximum and minimum stroke angles within the average wingbeat, respectively. We estimated the angle-of-attack α as the angle between the wing surface and its wingbeat-induced velocity vector (Fig. 3A).

We computed the wingbeat-induced angular velocity vector ($\mathbf{\Omega}=[\dot{\phi}, \dot{\eta}, \dot{\theta}]$) by analytically taking the time derivatives of the Fourier-series function of each wingbeat kinematic angle. Here, $\dot{\phi}$, $\dot{\eta}$ and $\dot{\theta}$ are the stroke rate, deviation rate and rotation rate, respectively. From these, we estimated the angular speed of the beating wings as $\omega=\sqrt{(\dot{\phi}^2+\dot{\eta}^2)}$.

We quantified the body kinematics during each wingbeat using the wingbeat-average flight speed, climb angle, and body pitch angle. For this, we first determined the temporal dynamics of flight velocity $\mathbf{U}=[u,v,w]$ as the temporal derivative of body positions. We then estimated the mean flight speed $U$ as the wingbeat-average magnitude of the flight velocity, and the corresponding climb angle as $\gamma_{climb}=\mathrm{atan}(U_{ver}/U_{hor})$. Here, $U_{ver}$ and $U_{hor}$ are the wingbeat-average vertical and horizontal components of flight velocity. The body pitch angle β was calculated as the angle between the body axis and the horizontal.

We assessed to what extent the studied wingbeats were representative of hovering flight by computing the advance ratio of all wingbeats as $J=U/(\bar{\omega}\ b)$, where $\bar{\omega}$ is the wingbeat-average angular wing speed. Hovering is generally defined as flight with advance ratios $J<0.1$, meaning that forward flight speed is less than 10% of the wingbeat-induced speed of the wingtip [20].

Because variability in flight measurements is typically high, especially compared to morphological measurements, we assessed the robustness of species-level differences in flight traits. For this, we compared within-species and between-species variation in each parameter to verify that our experiment effectively captured differences in flight kinematics between species. We applied a nested ANOVA model in which flight-sequence replicate was nested within species as a random effect. Comparison of sums of squares allowed assessment of variance attributable to differences between species versus within species. To facilitate visual comparison of interspecific versus intraspecific variation across parameters, we also computed the coefficient of variation for each parameter as $C_{var}$=(standard deviation/mean)·100% (Fig. S2).

#### *S1.5. Computational Fluid Dynamics simulations*

To estimate the aerodynamic forces and torques generated during hovering flight of the studied Diptera species, we performed Computational Fluid Dynamics (CFD) simulations. For each simulation, we modelled a single rigid, flat wing, using species-specific contours and wingbeat kinematics. The computational domain measured $10b \times 10b \times 10b$, and three complete wingbeat cycles were simulated. The wing thickness was set to $0.025b$. Assuming hovering flight conditions, we did not impose any mean inflow into the domain ($U_\infty$ = 0 m/s).

The simulations were carried out using our in-house open-source code WABBIT (Wavelet Adaptive Block-Based solver for Insects in Turbulence)[21,22]. This computational framework utilizes explicit finite-difference methods to solve the incompressible Navier–Stokes equations under an artificial-compressibility approach. Dynamic grid adaptation is achieved through wavelets, enabling local refinement of the discretization grid based on the flow at each time step. This ensures computational resources are allocated efficiently, with higher refinement where needed and coarser grids in regions of lesser importance. Starting from a coarse base grid, we allowed up to seven refinement levels, with each level doubling the grid's resolution. A Cohen–Daubechies–Feauveau wavelet (CDF60) was employed, and each block in the grid contained $B_s$ = 24 points per spatial direction, leading to an effective maximum resolution of 307 points per wingspan *b*. An additional simulation using eight refinement levels verified the accuracy of the results. As shown in prior validation tests, the numerical results are considered quasi-exact, equivalent to a physical experiment with the same rigid geometry and kinematics. For further details on the code, see Engels et al. (2021)[21]. The necessary parameter files to replicate these simulations are available in the online repository (see Data Availability Statement for details). To optimize performance, each simulation used up to 600 CPU cores and up to 2.7 TB of memory.

#### *S1.6. Modelling the aerodynamics and aeroacoustics of Diptera flight*

To interpret how morphology and kinematics shape aerodynamic force and power production and aero-acoustic output across species, we combined our full-DNS CFD simulations with a dimensional scaling framework.

**Modelling the aerodynamic forces produced by flapping insect wings**

We modelled the aerodynamic force production by flapping wings using a combined dynamic scaling and aerodynamic modelling approach. First, we used Buckingham-Pi theory [23] to determine the minimal set of dimensional parameters for modelling aerodynamic force production of an oscillating wing as

$$F \sim \rho\, b^4 f^2, \qquad \text{(S1)}$$

where ρ is air density, *b* is wingspan and *f* is the wingbeat frequency of the oscillating wing. First-order aerodynamic modelling states that the wingbeat-average aerodynamic thrust force produced during a wingbeat (*F*) scale with the product of dynamic pressure (*q*) and wing surface area (*S*) as *F~q S* [20]. Based on this, we model the aerodynamic thrust force produced by an insect wing as

$$F = \frac{1}{2}\rho\, S\, U^2\, C_F. \qquad \text{(S2)}$$

The aerodynamic force thus varies linearly with air density (ρ), wing area (*S*) and the thrust force coefficient ($C_F$), and quadratically with the wing speed *U*. During hovering flight, the effective wing speed can be approximated as the speed at the wings radius-of-gyration as $U_r = r_2\,\omega$, where ω is the angular velocity of the wing around the wing hinge, and $r_2$ is the wing radius-of-gyration relative to the wing hinge. A flying insect can vary this angular velocity by changing both its wingbeat frequency and amplitude, as angular velocity scales with the product of wingstroke amplitude and wingbeat frequency ($\omega \sim A_\phi f$). Thus, the aerodynamic force produced by an insect wing during hovering flight is

$$F = \frac{1}{2}\rho\, S\, {U_r}^2\, C_F = \frac{1}{2}\rho\, S\, (r_2\omega)^2\, C_F = \frac{1}{2}\rho\, S\, \left(r_2 f A_\phi\right)^2 C_F. \qquad \text{(S3)}$$

Finally, we converted this aerodynamic model to include only the minimal set of dimensional parameters of the Buckingham-Pi analysis (equation S8). For this, we decomposed wing area as $S = b\,\bar{c} = b^2/\mathrm{AR}$, where $\bar{c}$ is the mean wing chord, and AR is the wing aspect ratio. The radius-of-gyration can be decomposed as $r_2 = r_2^* \, b$, where $r_2^*$ is the span-normalised radius-of-gyration. As a result, the minimal model for estimating the effect of wing morphology, kinematics and aerodynamics on force production by two insect wings during hovering flight is

$$F = \frac{1}{\mathrm{AR}} \, \rho \, b^2 \, {U_r}^2 \, \mathrm{C}_F = \frac{1}{\mathrm{AR}} \, \rho \, b^4 \, (r_2^* \, \omega)^2 \, \mathrm{C}_F = \frac{1}{\mathrm{AR}} \, \rho \, b^4 \, (r_2^* \, A_\phi \, f)^2 \, \mathrm{C}_F. \tag{S4}$$

We validated this scaling against CFD-derived force estimations and then used it to interpret how species of different sizes produce weight support during hovering flight.

**Aerodynamic power from CFD and modelling**

We computed the instantaneous aerodynamic power directly from the CFD solutions as the dot product between the aerodynamic torque on a single wing and its angular velocity, summed over both wings and averaged over the wingbeat cycle. Specifically,

$$P_{\mathrm{aero}}(t) = 2 \, \mathbf{T}_{\mathrm{aero}}(t) \cdot \mathbf{\Omega}(t), \tag{S5}$$

where $\mathbf{T}_{\mathrm{aero}}(t)$ and $\mathbf{\Omega}(t)$ are the aerodynamic torque and angular-velocity vectors of the simulated wing, respectively. From this we determined the wingbeat-averaged power as the averaged over the last simulated wingbeat cycle, and reported its weight-normalized form, the weight-normalized aerodynamic power as

$$P_{\mathrm{aero}}^* = \frac{\bar{P}_{\mathrm{aero}}}{mg}, \tag{S6}$$

where *mg* is body weight. These CFD-derived quantities inherently include unsteady, viscous, and induced aerodynamic contributions.

For interpretation in the main text, we additionally refer to a compact, dimensionally consistent surrogate that separates kinematic and aerodynamic contributions as

$$P_{\mathrm{aero}}^* \propto U_D \, D^*, \tag{S7}$$

where $U_D \sim r_D \, f \, A_\phi$ is a characteristic wing speed at the drag moment arm $r_D$, and $D^* = 2D/mg = C_D/C_L$ is the weight-normalized wingbeat-average drag on the wing (equal to the inverse of the lift-to-drag ratio $C_L/C_D$). This surrogate is used only for explanatory scaling trends; all aerodynamic data plotted or tested statistically are computed directly from CFD. From CFD and kinematics we also estimated the normalized drag moment arm $r_D^* = r_D/b$ for all studied species, and compared this with the equivalent morphological metric, the normalized radius-of-gyration $r_2^*$ (Supplementary Fig. S3).

**Acoustic power from CFD-based aerodynamic forces**

Following the compact dipole description of aero-acoustic sources, far-field acoustic power is governed primarily by fluctuations of the aerodynamic pressure-force on the wing [24]. Therefore, the far-field acoustic power produced by beating wing can be estimated from CFD as

$$P_{\text{acoustic}} = \frac{1}{4\pi\rho c_s^3}\left(\overline{\frac{\partial F_p}{\partial t}}\right)^2, \tag{S8}$$

where $F_p$ is the air pressure force on the wing, and $c_s$ the speed of sound in air. Dividing and multiplying the acoustic power with wingbeat-averaged aerodynamic force $F$ leads to

$$P_{\text{acoustic}} = \frac{1}{4\pi\rho c_s^3}\left(\frac{\partial F_p}{\partial t}\right)^2 \frac{F^2}{F^2}. \tag{S9}$$

In hovering flight, the net force magnitude of both wings combined equals body weight ($2F = mg$). Because the net aerodynamic force and pressure force scale similarly with morphology and kinematics, the acoustic power can be rewritten as

$$P_{\text{acoustic}} = \frac{1}{4\pi\rho c_s^3}\left(\frac{\partial C_p}{\partial t^*}\frac{1}{C_F}\right)^2 (mgf)^2, \tag{S10}$$

Where normalized time equals $t^* = ft$, and $C_p$ and $C_F$ are the pressure and force coefficients, respectively. Based on this, we define the weight-normalized acoustic power as

$$P^*_{\text{acoustic}} = \frac{4\pi\rho c_s^3}{(mgf)^2} P_{\text{acoustic}} = \left(\frac{\partial C_p}{\partial t^*}\frac{1}{C_F}\right)^2, \tag{S11}$$

which separates the contribution of pressure-coefficient fluctuations $\partial C_p/\partial t^*$ from the mean force coefficient $C_F$ required for weight support. This normalization allows size-aware comparison of aero-acoustic output independent of the absolute mass and wingbeat frequency.

#### *S1.7. Mass-specific scaling of morphology, kinematics and aerodynamics under geometric, kinematic and dynamic similarity*

We used a dimensional similarity framework to quantify how wing morphology, wingbeat kinematics and aerodynamic forces scale with body mass under geometric, kinematic and dynamic similarity, respectively.

**<u>Mass-specific scaling of morphological parameters under geometric similarity</u>**

Under geometric similarity the morphological parameters should scale with body mass as following:

- The **wingspan *b*, mean wing chord $\bar{c}$** and **radius-of-gyration $r_2$** are length scales. Under geometric similarity, mass scales with length to the third power. Thus, wingspan, mean chord length and radius-of-gyration scale as $b \propto m^{1/3}$, $\bar{c} \propto m^{1/3}$, and $r_2 \propto m^{1/3}$ respectively.

- The **wing area *S*** is equal to the product of wingspan and mean chord ($S = b\bar{c}$). Substituting the scaling relationships for wingspan and mean chord into this equation gives $S \propto m^{1/3} \cdot m^{1/3} \propto m^{2/3}$.

- The **normalised radius-of-gyration $r_2^*$** and **aspect ratio AR** are wing shape parameters, which are independent of size. Thus, under geometric similarity both do not scale with mass as $r_2^* \propto m^0$ and AR $\propto m^0$.

###### Mass-specific scaling of wingbeat kinematics parameters under kinematic similarity

Under kinematic similarity, the wingbeat kinematic parameters **angular speed ω, wingbeat frequency $f$, angular amplitude $A_\Phi$**, and **angle-of-attack α** remain constant across sizes, and thus should not scale with body mass as $\omega \propto m^0$; $f \propto m^0$; $A_\Phi \propto m^0$; $\alpha \propto m^0$. The wingbeat kinematic parameter **wing speed $U_r$** scales with both wing kinematics and morphology as $U_r \propto f\, A_\Phi\, r_2$, and consequently scales with body mass under both kinematic and geometric similarity as $U_r \propto m^0\, m^0\, m^{1/3} = m^{1/3}$.

###### Mass-specific scaling of aerodynamic parameters under dynamic similarity

Under dynamic similarity, the aerodynamic force coefficient $C_F$ should remain constant, and thus should not scale with body mass as $C_F \propto m^0$.

#### *S1.8. Allometric scaling of morphology, kinematics and aerodynamics with mass for maintaining weight support across sizes*

For weight support during hovering flight, the upward-directed aerodynamic force produced by two wings should balance the weight of the animal ($2F = mg$). Thus, the aerodynamic force should scale linearly with body mass $F \propto m$. Furthermore, our aerodynamic model states that this force scales with wing morphology, kinematics and aerodynamics as in equation (S4). Based on these scaling laws we can estimate how a specific independent parameter should scale with mass, given that all other parameters scale with body mass under geometric, kinematic and dynamic similarity.

###### Allometric scaling of wing morphology with mass for maintaining weight support across sizes

The aerodynamic force scales with the wing morphology parameters $b$, $r_2^*$ and AR. Here, we determine how these morphological parameters should scale allometric for weight support, given that all other parameters scale with body mass under geometric, kinematic and dynamic similarity.

- The expected scaling of **wingspan *b*** for weight support, when all other parameters scale under geometric, kinematic and dynamic similarity, can be obtained by isolating $b$ in equation (S4), and substituting each element with its scaling relationships as

$$F \propto b^4\, r_2^{*2}/\mathrm{AR}$$

$$m \propto b^4\, m^0/m^0$$

$$b^4 \propto m^1$$

$$b \propto m^{1/4}.$$

Thus, to maintain weight support across sizes via allometric scaling of wingspan only, wingspan should scale with body mass as $b \propto m^{1/4}$.

- The expected scaling of the **normalized radius-of-gyration $r_2^*$** for weight support, given that all other parameters scale under geometric, kinematic and dynamic similarity, can be obtained by isolating $r_2^*$ in equation (S4), and substituting each element with its scaling relationships as

$F \propto b^4\, r_2^{*2}/\mathrm{AR}$

$m \propto m^{4/3}\, r_2^{*2}/m^0$

$r_2^{*2} \propto m^{1+0-4/3} = m^{-1/3}$

$r_2^* \propto m^{-1/6}$.

Thus, to maintain weight support via allometric scaling of the span-normalized radius-of-gyration only, this should scale with body mass as $r_2^* \propto m^{-1/6}$.

- The expected scaling of the wing's **aspect ratio AR** for weight support, given that all other parameters scale under geometric, kinematic and dynamic similarity, can be obtained by isolating AR in equation (S4), and substituting each element with its scaling relationships as

$F \propto b^4\, r_2^{*2}/\mathrm{AR}$

$m \propto m^{4/3}\, m^0/\mathrm{AR}$

$\mathrm{AR} \propto m^{4/3+0-1} = m^{1/3}$.

Thus, to maintain weight support via allometric scaling of aspect ratio only, this should scale with body mass as $\mathrm{AR} \propto m^{1/3}$.

**<u>Allometric scaling of wingbeat kinematics with mass for maintaining weight support</u>**

The aerodynamic force scales with the wingbeat kinematics parameters $U_r$, $\omega$, $f$ and $A_\phi$. Here, we determine how these kinematics parameters should scale allometric with body for maintaining weight support, given that all other parameters scale with body mass under geometric, kinematic and dynamic similarity.

- The expected scaling of the **wing speed $U_r$** with body mass for weight support, when all other parameters scale under geometric, kinematic and dynamic similarity, can be obtained by isolating $U_r$ in equation (S4) and substituting each element with its specific mass scaling relationship as

$F \propto b^2\, (U_r)^2$

$F \propto b^2\, {U_r}^2$

$m \propto m^{2/3} \cdot {U_r}^2$

${U_r}^2 \propto m^{1-2/3} \propto m^{1/3}$

$U_r \propto m^{1/6}$.

Thus, to maintain weight support across body masses via allometric scaling of the angular wing speed only, it should scale with body mass as $\omega \propto m^{-1/6}$.

- The expected scaling of the **wing angular speed ω** with body mass for weight support, when all other parameters scale under geometric, kinematic and dynamic similarity, can be obtained by isolating ω in equation (S4) and substituting each element with its specific mass scaling relationship as

$$F \propto b^4\, \omega^2$$

$$m \propto m^{4/3} \cdot \omega^2$$

$$\omega^2 \propto m^{1-4/3} \propto m^{-1/3}$$

$$\omega \propto m^{-1/6}.$$

Thus, to maintain weight support across body masses via allometric scaling of the angular wing speed only, it should scale with body mass as $\omega \propto m^{-1/6}$.

- The expected scaling of the sub-components of wing speed **wingbeat frequency *f*** and **stroke amplitude $A_\phi$** for weight support, when all other parameters scale under geometric, kinematic and dynamic similarity, are the same as that of angular wing speed itself as

$$\omega \propto f\, A_\phi$$

$$A_\phi \propto m^0 \;\rightarrow\; f \propto m^{-1/6}$$

$$f \propto m^0 \;\rightarrow\; A_\phi \propto m^{-1/6}.$$

Thus, to maintain weight support across body masses via allometric scaling of wingbeat frequency or amplitude only, either of them should scale with body mass as $f \propto m^{-1/6}$ or $A_\phi \propto m^{-1/6}$.

**Allometric scaling of aerodynamic forces with mass for maintaining weight support**

The aerodynamic force scales linearly with the aerodynamics parameter $C_F$. The expected scaling of the **force coefficient $C_F$** with body mass for weight support, when all other parameters scale under geometric, kinematic and dynamic similarity, can be obtained by isolating $C_F$ in equation (S4) and substituting each element with its specific mass scaling relationship as

$$F \propto b^4\, C_F$$

$$m \propto m^{4/3}\, C_F$$

$$C_F \propto m^{1-4/3}$$

$$C_F \propto m^{-1/3}.$$

Thus, to maintain weight support across body masses via allometric scaling of the aerodynamic force coefficient only, it should scale with body mass as $C_F \propto m^{-1/3}$.

**Mass-specific scaling of aerodynamic power under similarity, and for maintaining weight support**

As described in section S1.6, the weight-normalized power scales as $P^*_{\text{aero}} = U_r\ D^* = U_r\ C_D/C_L$, where $C_L$ and $C_D$ are the lift and drag force coefficients, respectively. Based on this scaling law, we determined the scaling of weight-normalized power with body mass, under geometric, kinematic and dynamic similarity, and for maintaining weight support (2*F=mg*):

1. Under dynamic similarity, the lift and drag force coefficient do not scale with body mass ($\propto m^0$); under morphological and kinematic similarity wing speed scales as $U_r \propto m^{1/3}$. As a result, under geometric, kinematic and dynamic similarity weight-normalized power should scale with mass as $P^*_{\text{aero}} \propto m^{1/3}$.
2. To maintain weight support through variations in wing speed only (dynamic similarity: $D^* \propto m^0$), wing speed should scale with body mass as $U_r \propto m^{1/6}$. Consequently, when maintaining weight support across sizes, weight-normalized power should also scale with mass as $P^*_{\text{aero}} \propto m^{1/6}$.

**Mass-specific scaling of aero-acoustic sound power under similarity, and for maintaining weight support**

As described above in S1.6, acoustic power ($P_{\text{acoustic}}$) and the corresponding normalized acoustic-power ($P^*_{\text{acoustic}}$) produced by a flapping wing scales as defined in Equations (S10) and (S11), respectively. Based on these scaling laws, we determined the scaling of (normalized) acoustic power with body mass, under geometric, kinematic and dynamic similarity, and for maintaining weight support (*F=mg*):

1. Under geometric, kinematic and dynamic similarity, all parameters of interest do not scale with mass ($\frac{\partial \text{C}_\text{p}}{\partial \text{t}^*} \propto m^0$; $C_F \propto m^0$; $f \propto m^0$), and thus the acoustic power scale with mass as $P_{\text{acoustic}} \propto m^2$. The corresponding weight-normalized acoustic power does not scale with mass ($P^*_{\text{acoustic}} \propto m^0$).
2. To maintain weight support through variations in wingbeat frequency alone, wingbeat frequency should scale with body mass as $f \propto m^{-1/6}$. Consequently, when maintaining weight support across sizes under dynamic similarity ($\frac{\partial \text{C}_\text{p}}{\partial \text{t}^*} \propto m^0$; $C_F \propto m^0$), the acoustic power scale with mass as $P_{\text{acoustic}} \propto m^2 m^{-1/3} \propto m^{5/3}$. The corresponding weight-normalized acoustic power does not scale with mass ($P^*_{\text{acoustic}} \propto m^0$).

#### *S1.9. Allometric adaptations and weight-support contributions*

To quantify how individual traits contribute to maintaining weight support across body size, we compared their observed allometric slopes with two theoretical expectations: (i) the slope predicted under geometric, kinematic, and dynamic similarity ($a_{\text{sim}}$; see section S1.7), and (ii) the slope required for maintaining weight support if that trait alone were adjusted ($a_{\text{ws}}$; see section S1.8). To do so, we computed the relative allometric scaling for weight support for each trait as [25]

$$a^* = \frac{a_{\text{allo}} - a_{\text{sim}}}{a_{\text{ws}} - a_{\text{sim}}} \times 100\%, \tag{S12}$$

where $a_{\text{allo}}$ is the allometric PGLS slope fitted to the experimental or CFD data. This metric expresses how strongly the observed scaling of a trait shifts away from similarity and toward the theoretical requirement for maintaining weight support when considered in isolation. Positive values indicate movement toward the weight-support expectation, whereas negative values indicate movement in the opposite direction. This approach provides a consistent basis for comparing the relative importance of morphological, kinematic, and aerodynamic traits in supporting body weight during hovering [25].

#### *Supplementary references*

7. Bookstein F. L.. Morphometric tools for landmark data geometry and biology. Cambridge University Press; 1991.

24. Lighthill James. Waves in fluids. Cambridge University Press; 2001.

25. Le Roy C, Tervelde N, Engels T, Muijres FT. Adaptations in wing morphology rather than wingbeat kinematics enable flight in small hoverfly species. Elife. 2025;13. doi:10.7554/eLife.97839
