## Supplementary figures and images for "Diptera flight diversity is shaped by aerodynamic constraints, scaling, and evolutionary trade-offs"

### S1 Fig.

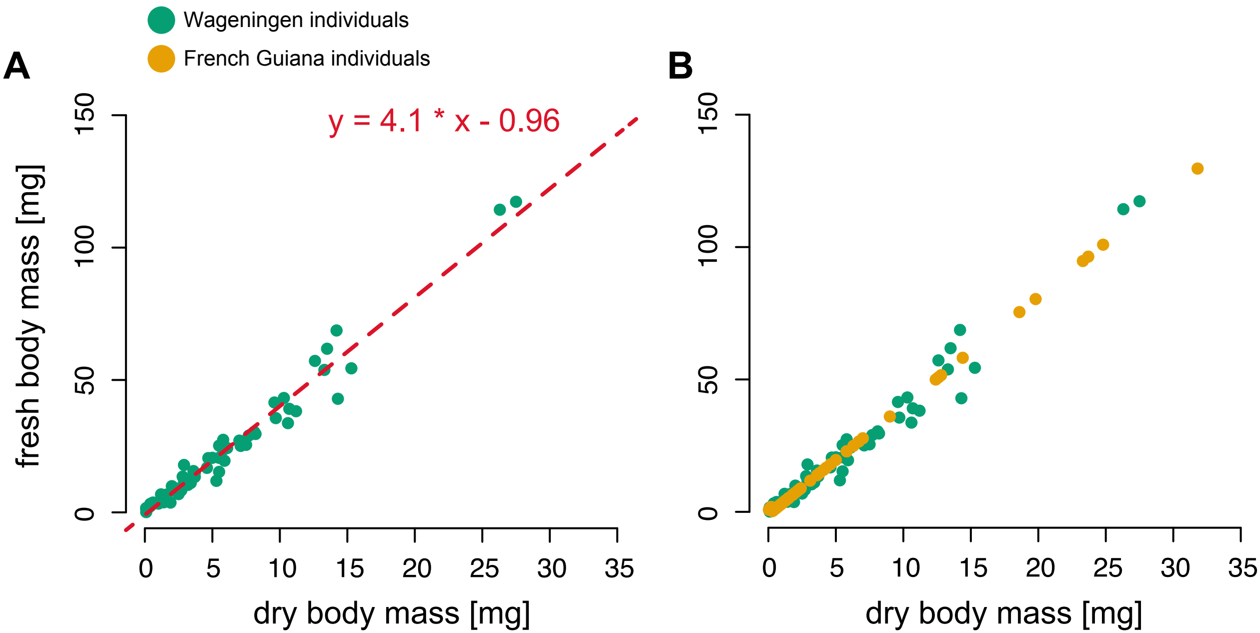

### S1 Table

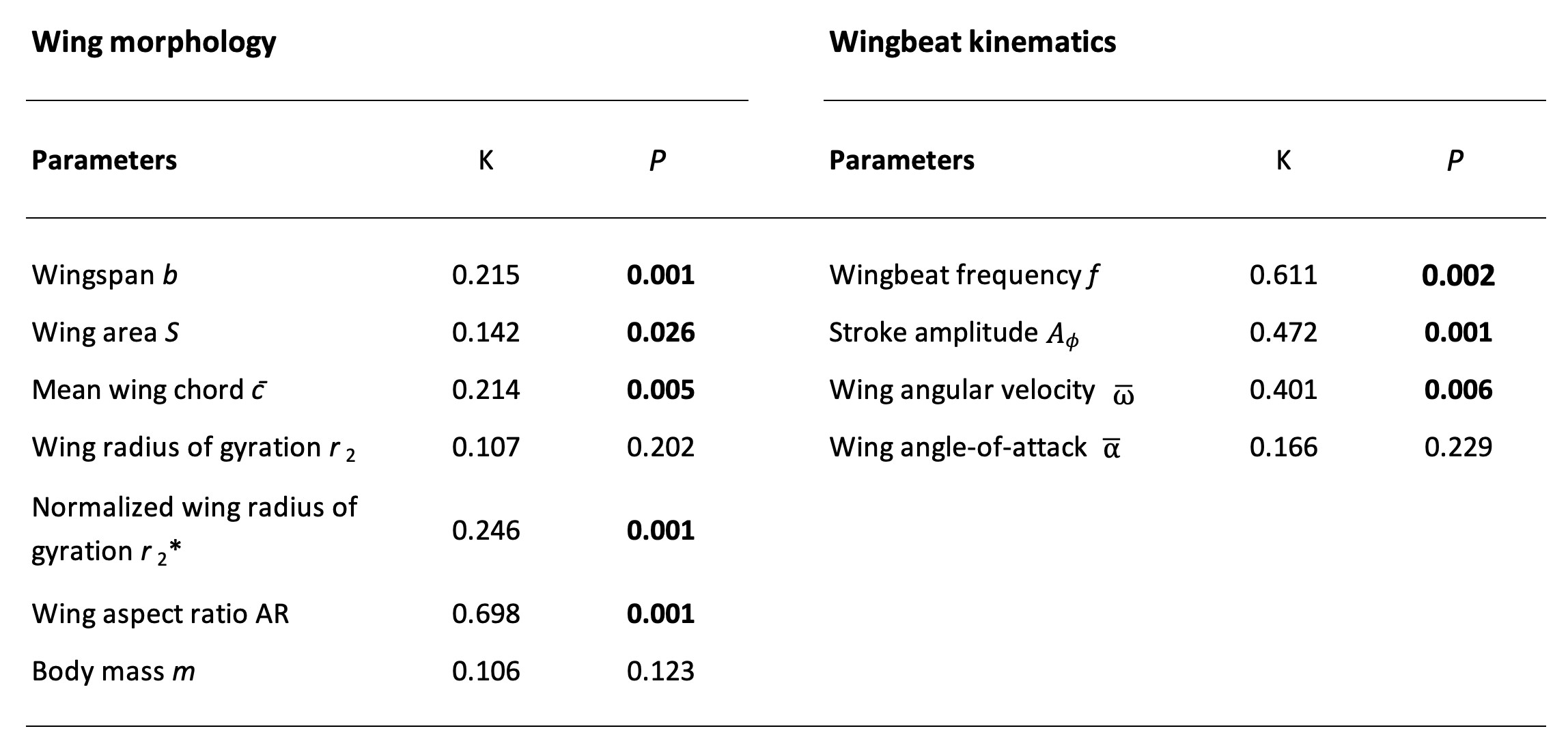

### S2 Fig.

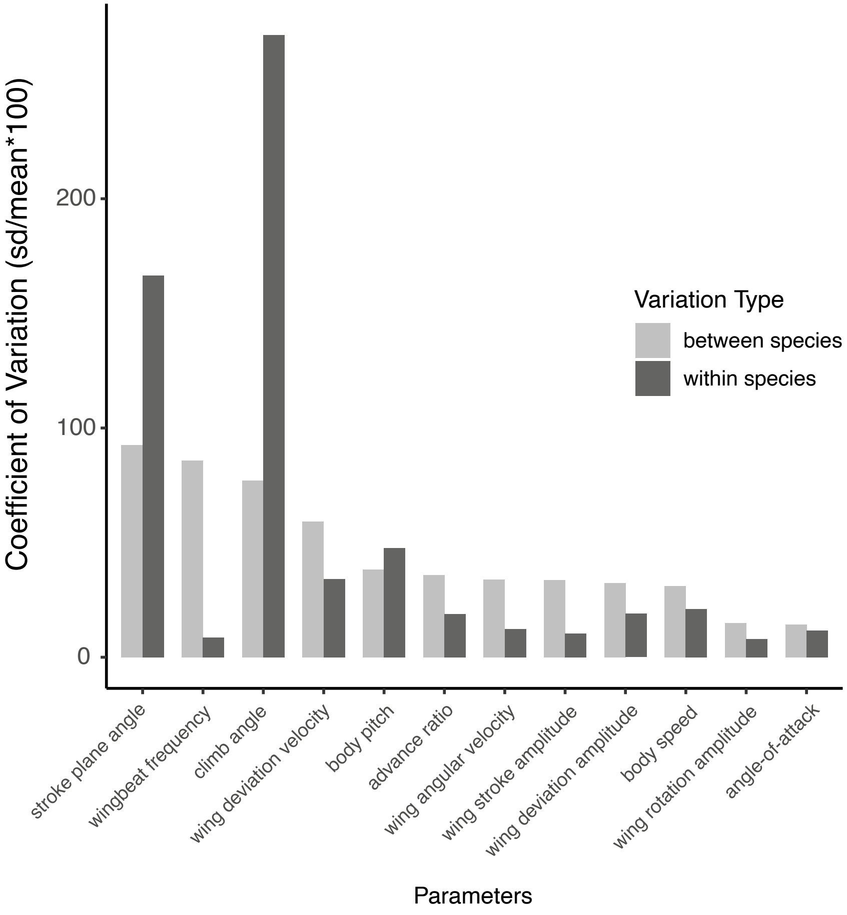

### S2 Table

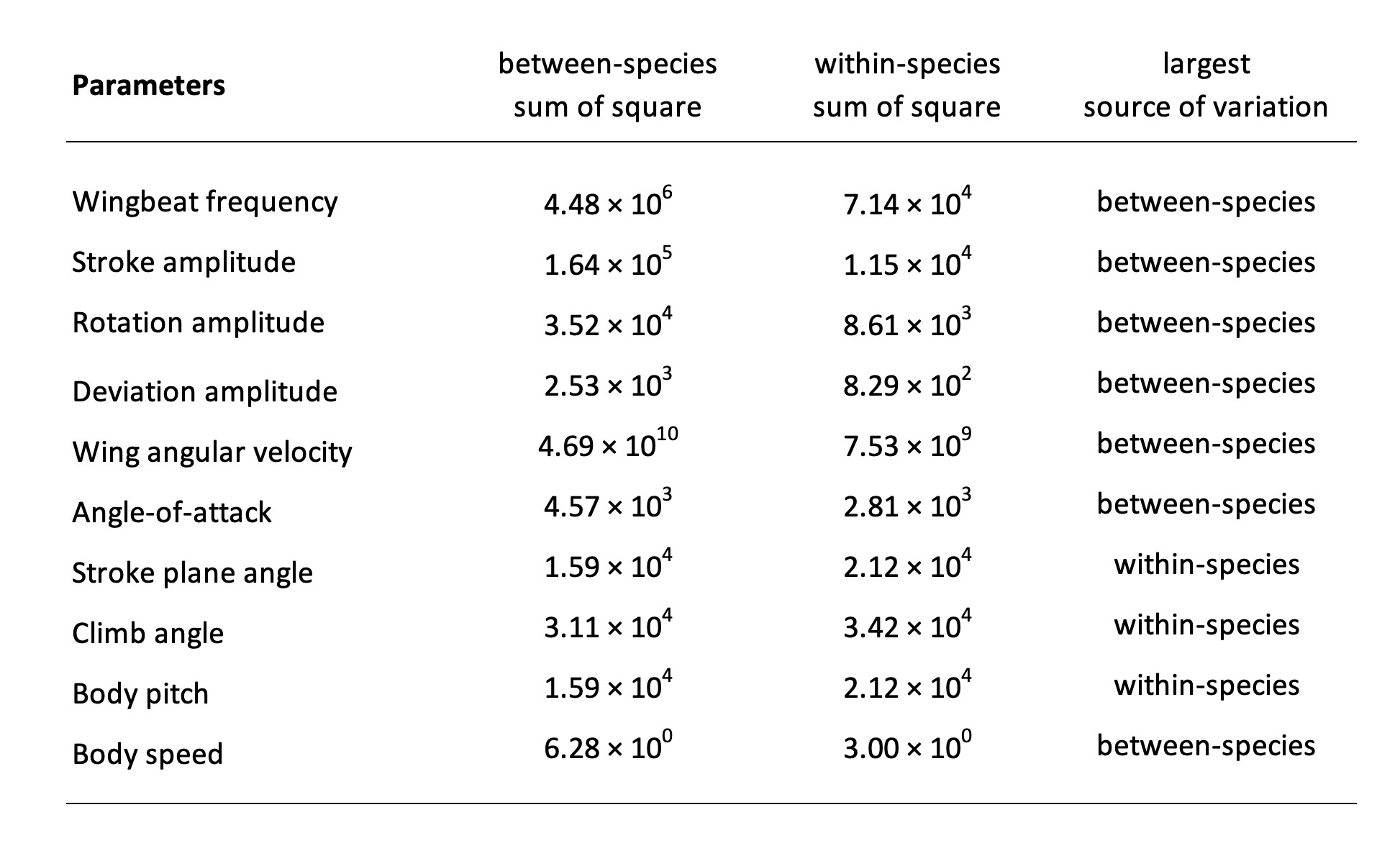

### S3 Fig.

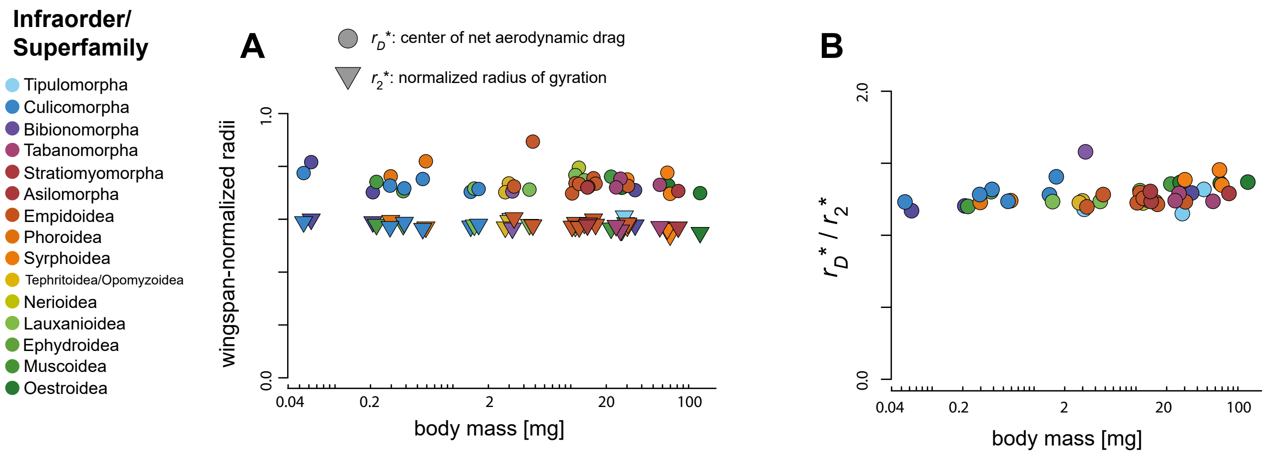
